# Gastrointestinal colonization as a source of *Staphylococcus aureus* in atopic dermatitis

**DOI:** 10.1101/2025.04.17.648849

**Authors:** Theodora K. Karagounis, Greg Putzel, Magdalena Podkowik, Julia Shenderovich, Alice Tillman, Sabrina M. Morales, Abigail Gaylord, Anusha Srivastava, Natalia Arguelles, Hannah Slater, Abonti N. Ahmed, Yue Xing, Nora M. Samhadaneh, Carli D. Needle, Robert J. Ulrich, Vikash Oza, Jonas Schluter, Shruti Naik, Leonardo Trasande, Alejandro Pironti, Victor J. Torres, Bo Shopsin

**Author notes:** **Co-Corresponding authors:** Theodora Karagounis MD, MS; Alexandria Center for Life Science West Tower, 430 E 29^th^ St, New York, NY 10016, Bo Shopsin MD, PhD; Alexandria Center for Life Science West Tower, 430 E 29^th^ St, New York, NY 10016.

## Abstract

Atopic dermatitis (AD) is a prevalent inflammatory skin disease with complex pathogenesis. Both skin and gut microbiota influence AD, with *Staphylococcus aureus*, in particular, exacerbating the disease. However, the relationship between *S. aureus* colonization in the gut and skin, and whether it affects AD, remains unclear. Using a combination of culture-based methods, microbiome analysis, and genome sequencing of *S. aureus* from multiple body sites of children with and without AD, we found that the gut represents a major reservoir of genetically diverse *S. aureus* that is transmitted to the skin, including mutants associated with worse disease. We validated this association between *S. aureus* gastrointestinal colonization and AD in an independent human cohort and demonstrated its direct effect on disease in an infantile AD mouse model, wherein *S. aureus* gastrointestinal colonization worsened skin inflammation. Overall, this study identifies a previously unrecognized *S. aureus* reservoir, with implications for microbiota-targeting therapies in AD.

## INTRODUCTION

Atopic dermatitis (AD) affects up to 20% of children and classically starts in infancy with pruritic red patches of the skin followed later by the development of co-morbid allergic conditions.^1,2^ To-date, most AD therapies target locally or systemically dysregulated inflammatory pathways and require chronic treatment, sometimes with resultant side effects.^3^ These limitations in management options for AD highlight the need for a deeper understanding of its underlying mechanisms and discovery of alternative targets for disease treatment and prevention.

AD pathophysiology is multifactorial, shaped by host genetics and environmental exposures. However, these factors alone do not fully explain its widespread prevalence.^4,5^ Increasingly, microbial influences are recognized as critical components of the pathophysiology of inflammatory skin diseases,^6^ with AD serving as the prototypical example.^7–9^ The microbiota, especially *Staphylococcus aureus*, contribute to AD pathology,^6^ as the presence and density of *S. aureus* on the skin correlates with disease development and flare.^7,8,10^ While the nares have traditionally been considered the primary site of *S. aureus* colonization,^11^ gastrointestinal (GI) carriage —common in infants^12^— has been associated with an increased risk of *S. aureus* infection and is frequently observed in children with AD.^13,14^ Notably, as infants age and their gut microbiome matures, *S. aureus* GI colonization declines, coinciding with a decreasing age-dependent prevalence in infantile AD and suggesting a link between GI *S. aureus* and AD.^12^ However, despite these associations, the role of *S. aureus* gut colonization in promoting AD remains unclear.

*S. aureus* drives skin inflammation through toxins and proteases that activate mast cells^15^, break down the skin barrier^16^, and induce itch.^17^ These processes are upregulated by the *S. aureus agr* quorum-sensing system^18^, which plays a role in both GI and skin colonization.^19–21^ Paradoxically, however, *agr* loss-of-function mutations in *S. aureus* have been observed on the skin of AD subjects,^22^ suggesting that within-host evolution of *S. aureus* may contribute to AD pathogenesis. Understanding such adaptive changes could provide mechanistic insight and lead to novel therapeutic strategies that target bacterial pathways to better manage AD.

In the present study, we hypothesized that the GI tract serves as a reservoir of *S. aureus* that contributes to pediatric AD. By combining culture data and whole-genome sequencing of *S. aureus* from multiple body sites in a longitudinal cohort of children, we identified the GI tract as a disease-relevant reservoir from which *S. aureus* is transmitted to the skin, establishing a direct relationship between these two distal microbial niches. We validated this association in an independent cohort and identified key factors influencing *S. aureus* GI colonization, including adaptive bacterial mutations and supportive co-colonizers. Complementing our human data, we show that GI colonization by *S. aureus* drives AD pathogenesis in a mouse model of infantile AD. Together, these results support a novel way of conceiving the role of gut microbiota in inflammatory skin diseases – namely, as a bacterial reservoir and source of genetic diversity for skin-colonizing pathogens.

## RESULTS

### *S. aureus* gastrointestinal colonization is associated with pediatric AD

To determine whether *S. aureus* GI colonization is associated with pediatric AD, we conducted a prospective observational study of 83 children (ages 0.6-13.6 years; average 3.6 years), enrolled from 2021 to 2024. Of those, 61 subjects had AD and 22 were healthy controls (see **Fig. 1a** and for details **Extended Data Table 1** and **Methods**).

**Fig. 1:**
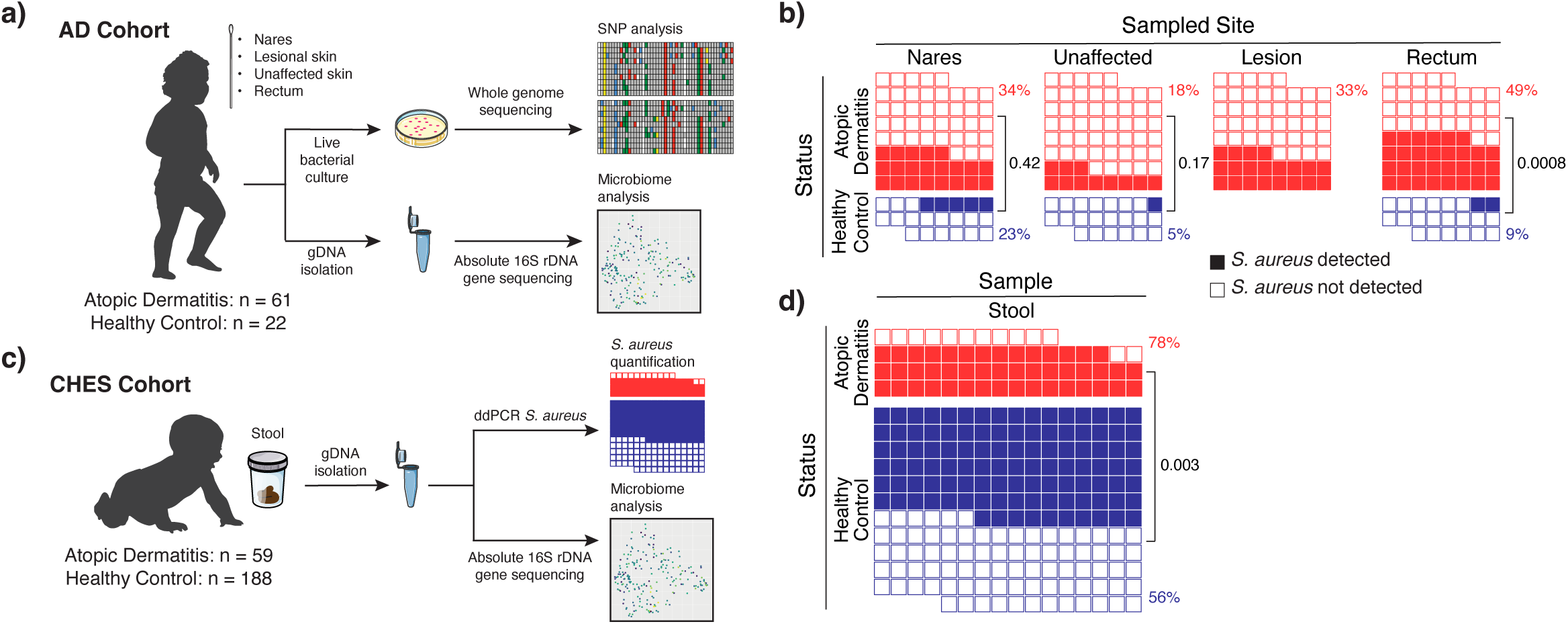
*S. aureus* gastrointestinal colonization is associated with atopic dermatitis. **a)** Overview of the AD cohort study design. **b)** Proportion of subjects colonized by *S. aureus* in the AD cohort. Percent of subjects with *S. aureus* cultured from the first timepoint at each sampled body site. Each box represents one subject; filled boxes indicate *S. aureus* detection, and unfilled boxes indicate no detection. **c)** Overview of the CHES cohort study design. **d)** Proportion of subjects colonized by *S. aureus* in the CHES cohort. Percent of CHES cohort subjects colonized with *S. aureus*, determined by droplet digital PCR targeting the *S. aureus*-specific *spa* gene. Statistical significance assessed by Fisher’s exact test.

To assess *S. aureus* colonization, we cultured bacteria from lesional and non-lesional skin, nares, and rectum—the latter serving as a proxy for lower GI colonization.^23,24^ Consistent with prior studies, approximately 30% of AD subjects were colonized on lesional skin (**Fig. 1b**), and 20% on non-lesional skin, compared to 5% in controls (**Fig. 1b**).^25^ While nares colonization rates were similar between AD subjects and controls (34% vs 23%, *p*=0.42; **Fig. 1b**), rectal colonization rates were more than 5-fold higher in AD subjects (49% vs 9%, *p*=0.0008; **Fig. 1b**), suggesting a specific association between GI colonization and AD. To validate the use of rectal swabs as a proxy for GI colonization, we analyzed 27 available matched stool samples from 19 AD patients for the presence of *S. aureus* using digital droplet (dd)PCR, given concerns about bacterial viability during mail-in transit to the lab. *S. aureus* was detected by ddPCR in all positive rectal cultures, confirming high specificity of culture of rectal swabs (area under receiver operating characteristic curve: 0.79, **Extended Data Fig. 1a**). ddPCR results highly correlated with culture results (*R*=0.8, *p*<0.0001, **Extended Data Fig. 1b**). In summary, *S. aureus* colonization is more common in the GI tracts of children with AD than in controls, whereas nasal colonization does not differ significantly between groups (**Fig. 1b**).

To validate and expand our findings, we applied the ddPCR assay to 352 stool lysates from a second human cohort, aged 6 to 14 months, enrolled in the New York University (NYU) Children’s Health and Environment Study (CHES, see **Fig. 1c**).^26^ This study, which is a prospective birth cohort, included 59 (24%) infants with AD and 188 healthy controls (infants without a record of AD). In this cohort, *S. aureus* was detected in 78% of infants with AD, compared to 56% in controls (*p*=0.003, **Fig. 1d**). The higher detection rate of *S. aureus* in the CHES cohort compared to the AD cohort likely reflects the increased sensitivity of PCR vs. culture methods,^27^ as well as the younger age of this cohort (average age 11 months in CHES and 46 months in the AD study). These data are consistent with prior reports that younger children are more frequently colonized with *S. aureus* in the GI tract than older children.^12^ Collectively, results from the CHES cohort, a robust independent prospective sample collection, are consistent with those of the AD cohort and support the conclusion that *S. aureus* GI colonization is associated with AD.

### *S. aureus* gastrointestinal colonization is associated with AD severity

In our AD cohort, most subjects had mild to moderate disease, consistent with the broader AD population (first timepoint severity: mild 39%, moderate 49%, and severe 12%).^28^ *S. aureus* skin lesion colonization was observed in 33% of AD subjects, concordant with prior studies on children with mild-to-moderate disease.^25^ At the rectal site, *S. aureus* presence and density correlated with disease severity. This pattern mirrors findings at the lesional skin site, suggesting a direct link between *S. aureus* GI colonization and AD severity (**Fig. 2a-e, Extended Data Table 2**).^25^

**Fig. 2:**
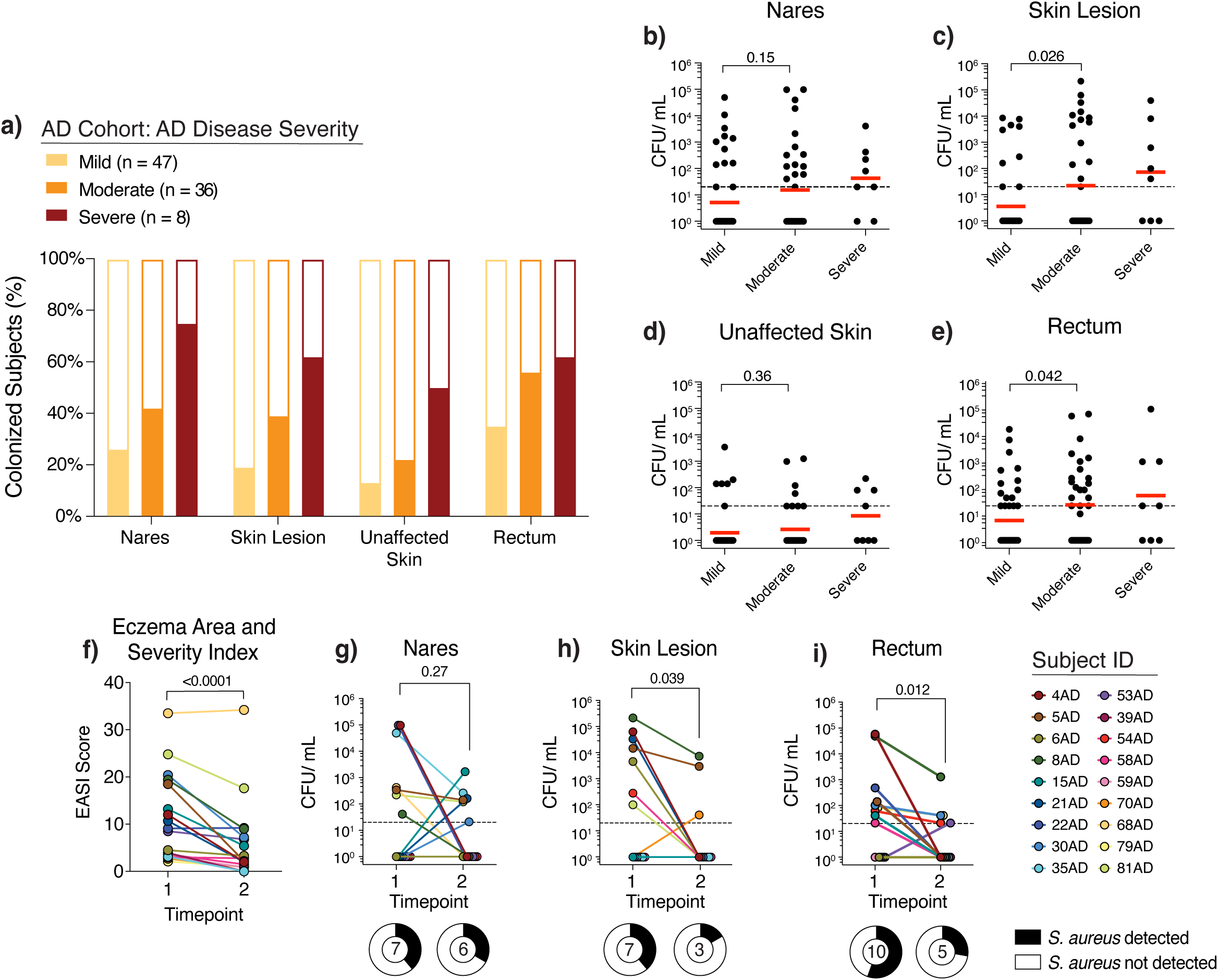
*S. aureus* gastrointestinal colonization mirrors lesional skin colonization and is associated with disease severity and flare. Disease severity was assessed using the Eczema Area and Severity Index (see **Methods**). **a)** Study subjects with *S. aureus* cultured at each body site, grouped by disease severity at the time of sampling. b-e) *S. aureus* colonization density by body site. Density in the **b)** nares, **c)** lesional skin, **d)** unaffected skin, and **e)** rectum, grouped by disease severity at the time of sampling. Red lines indicate the geometric mean while dotted lines indicate the limit of detection at 20 CFU/mL; significance tested by Mann-Whitney U Test. Significance testing not performed with the severe group due to underpowering of this group. **f)** AD severity over time. EASI score over time, indicating AD disease severity. g-i) *S. aureus* colonization density over time. Density over time in the **g)** nares, **h)** lesional skin, and **i)** rectum, based on culture results. Dashed lines indicate the detection limit of 20 CFU/mL. Pie charts depict the proportion of individuals with *S. aureus* detected or not detected at each time point, with the center number indicating number of individuals in whom *S. aureus* was detected. Statistical significance assessed by Wilcoxon matched pairs signed rank test.

Since AD fluctuates between periods of flare and remission, we evaluated available longitudinal samples from follow-up visits in 18 of 61 AD subjects. We observed that *S. aureus* colonization on lesional skin and in the rectum—but not in the nares—decreased with flare resolution by, on average, 100-fold (**Fig. 2f-i**), suggesting a reciprocal link between skin health and GI colonization. Notably, the reduction in GI colonization after flare resolution was not age-related (**Extended Data Fig. 2**), indicating that this change is directly associated with AD activity, rather than aging. Together, these findings show that *S. aureus* colonization dynamics in the gut temporally mirror skin colonization and track with AD flares, pointing to a functional role for GI-derived *S. aureus* in AD pathogenesis. Data on AD severity and flare timing for the CHES cohort were unavailable.

### GI tract colonization by *S. aureus* is associated with global gut microbiome shifts

Persistence of an immature microbiota, which lacks resistance to *S. aureus* colonization, has been linked to AD.^29^ To explore this phenomenon, we characterized the GI microbiome from rectal swabs of 49 of 83 AD cohort individuals using quantitative 16S rDNA sequencing.^30,31^

Gut microbiome diversity typically increases after birth and plateaus by age two years.^32^ With the majority of the AD cohort aged older than 2 years (67% > 2 years), we observed a weak relationship between age and alpha diversity (*p*=0.05, **Extended Data Fig. 3a**). Bray-Curtis dissimilarity, a beta-diversity measure, revealed age-related compositional differences (*p*=0.001, **Extended Data Fig. 3b**). These findings indicate the presence of age-related shifts in microbial composition despite limited overall diversity increases.

In addition to global, age-related shifts, microbiome diversity was reduced in *S. aureus* GI-colonized individuals. AD subjects with rectal *S. aureus* colonization had lower alpha diversity than non-colonized individuals (*p*=0.019, **Fig. 3a**). Considering all AD cohort subjects, Bray-Curtis dissimilarity differed significantly by *S. aureus* colonization status (*p*=0.039, **Fig. 3b**), but not by disease status or AD severity (**Fig. 3c, Extended Data Fig. 3c**). These results indicate that *S. aureus* presence in the gut, independent of AD, is directly linked to microbiome changes.

**Fig. 3:**
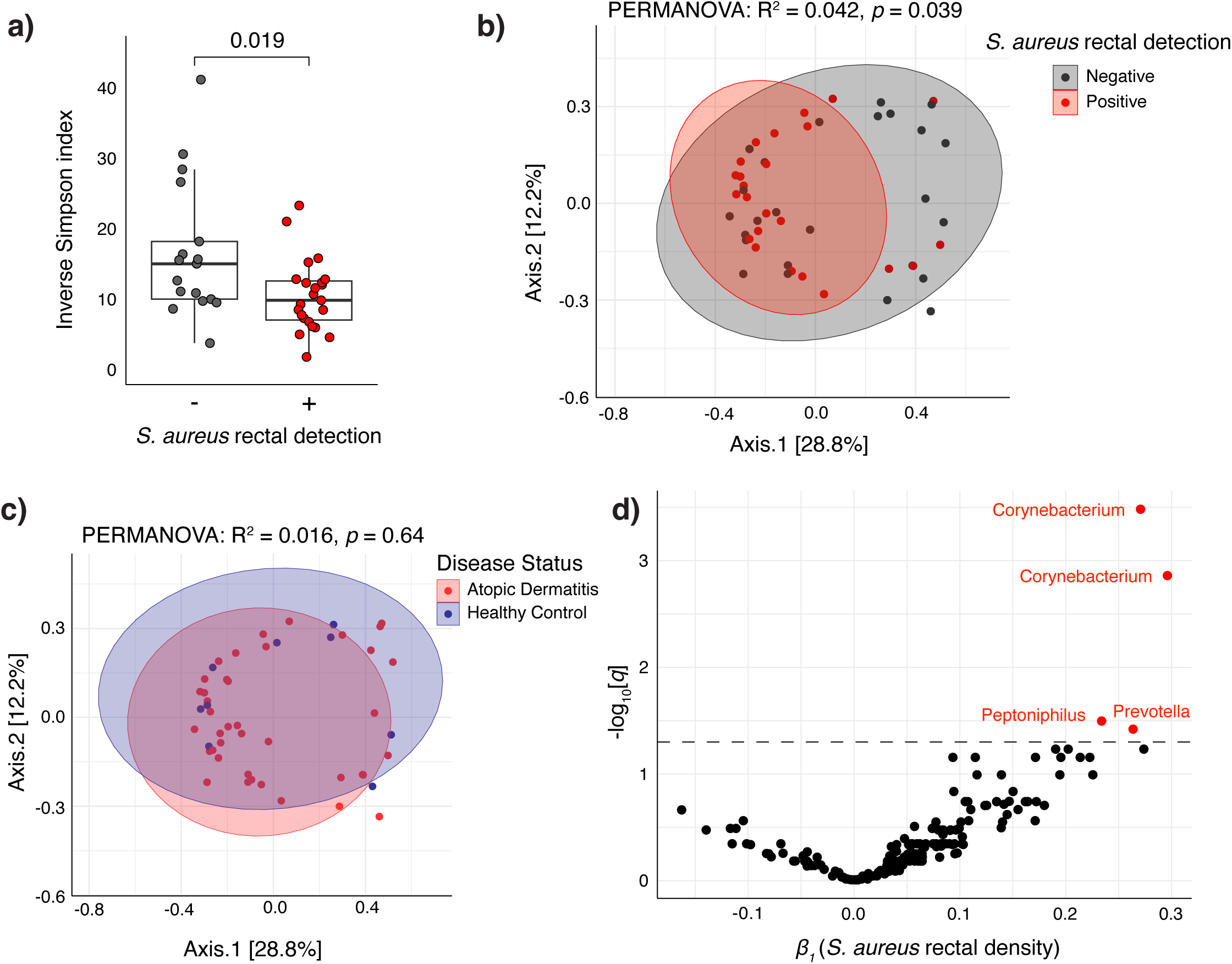
Gastrointestinal *S. aureus* colonization is associated with global gut microbiome changes in atopic dermatitis. **a)** Alpha diversity by *S. aureus* colonization in AD subjects. Comparison of the inverse Simpson index between AD subjects with and without *S. aureus* detected in rectal swabs by culture (n = 40 subjects). Statistical significance assessed by Mann-Whitney U test. **b-c)** Bray-Curtis dissimilarity, demonstrating community composition differences by **b)** *S. aureus* presence in rectal swabs by culture or **c)** disease status (n = 40 AD, 9 HC subjects). Statistical significance assessed by PERMANOVA. **d)** Differential abundance of amplicon sequence variants (ASVs) by *S. aureus* rectal density (as determined by culture). Volcano plot showing ASVs that are differentially abundant by *S. aureus* rectal density, modeled using a linear mixed effects model (see **Methods**). The x-axis indicates the *S. aureus* rectal density coefficient. The q-value indicates the significance of the *S. aureus* coefficient correlating with the absolute abundance of the indicated taxon with Benjamini–Hochberg correction for multiple hypothesis testing. The dotted line represents a 5% false-discovery rate threshold. Significant ASVs labelled by genus. Inverse Simpson index assessed at ASV level. Bray Curtis dissimilarity calculated at genus level.

To investigate differences in the density of specific bacterial taxa, we normalized the relative abundances of amplicon sequence variants (ASVs) to a bacterial spike-in (*Salinibacter ruber*) absent from the mammalian microbiome, as previously described.^30,31^ Among 183 ASVs with ≥10% prevalence, ASVs from the *Corynebacterium* genus and from *Prevotella corporis* and *Peptoniphilus coxii* species correlated positively with *S. aureus* rectal culture density (linear mixed-effects model; **Fig. 3d**), while no taxon was significantly negatively associated. Prior work investigating eczematous skin lesions in a murine AD model similarly observed co-colonization of *Corynebacterium* species and *S. aureus* suggesting these species may positively interact within this disease setting.^33^

We then examined the CHES cohort (ages 6-14 months). In this younger group, alpha and beta diversity varied strongly with age (*p*<0.001, *p*=0.001, respectively, **Extended Data Fig. 3d,e**), consistent with rapid early-life microbiome development.^32^ Unlike the older AD cohort, *S. aureus* colonization was not significantly associated with diversity or compositional differences though the latter approached significance (*p*=0.073) (**Extended Data Fig. 3f,g**). Consistent with age-related diversity trends that may mask *S. aureus*-associated effects, CHES cohort alpha diversity was lower than in the older AD cohort (*p=*0.001, with median ages 11.8 months and 37 months, respectively; **Extended Data Fig. 3h**).

### AD subjects are colonized with diverse *S. aureus* clones

The observed correlation between *S. aureus* GI colonization and AD severity and flare (**Fig. 2**) suggests that the GI tract may serve as a reservoir for seeding affected skin. To explore this possibility, we used bacterial whole-genome sequencing to analyze *S. aureus* strains isolated from skin, nasal, and rectal sites of patients in our AD cohort. Variation in the density and presence of *S. aureus* across colonization sites and disease severity resulted in differences in the number of isolates recovered across sites, subjects, and timepoints (**Extended Data Table 2**). For each body site, we aimed to sequence 10 single-colony isolates,^34^ resulting in whole-genome sequences from 38 AD patients and 7 healthy controls. This yielded a total number of 412, 407, and 338 isolates from the nares, skin, and rectum, respectively.

*S. aureus* lineages were defined to encompass all isolates pairs with fewer than 50 single-nucleotide variants (SNVs) across the core genome (see **Methods**). Using this threshold, most isolates within individuals were more closely related to each other than to isolates from other participants, suggesting that these strains originated from a common ancestor within the host rather than from mixed colonization by multiple circulating strains (**Fig. 4a**). Consistent with prior work, 8 of 38 (∼20%) subjects were colonized with more than one lineage.^35,36^

**Fig. 4:**
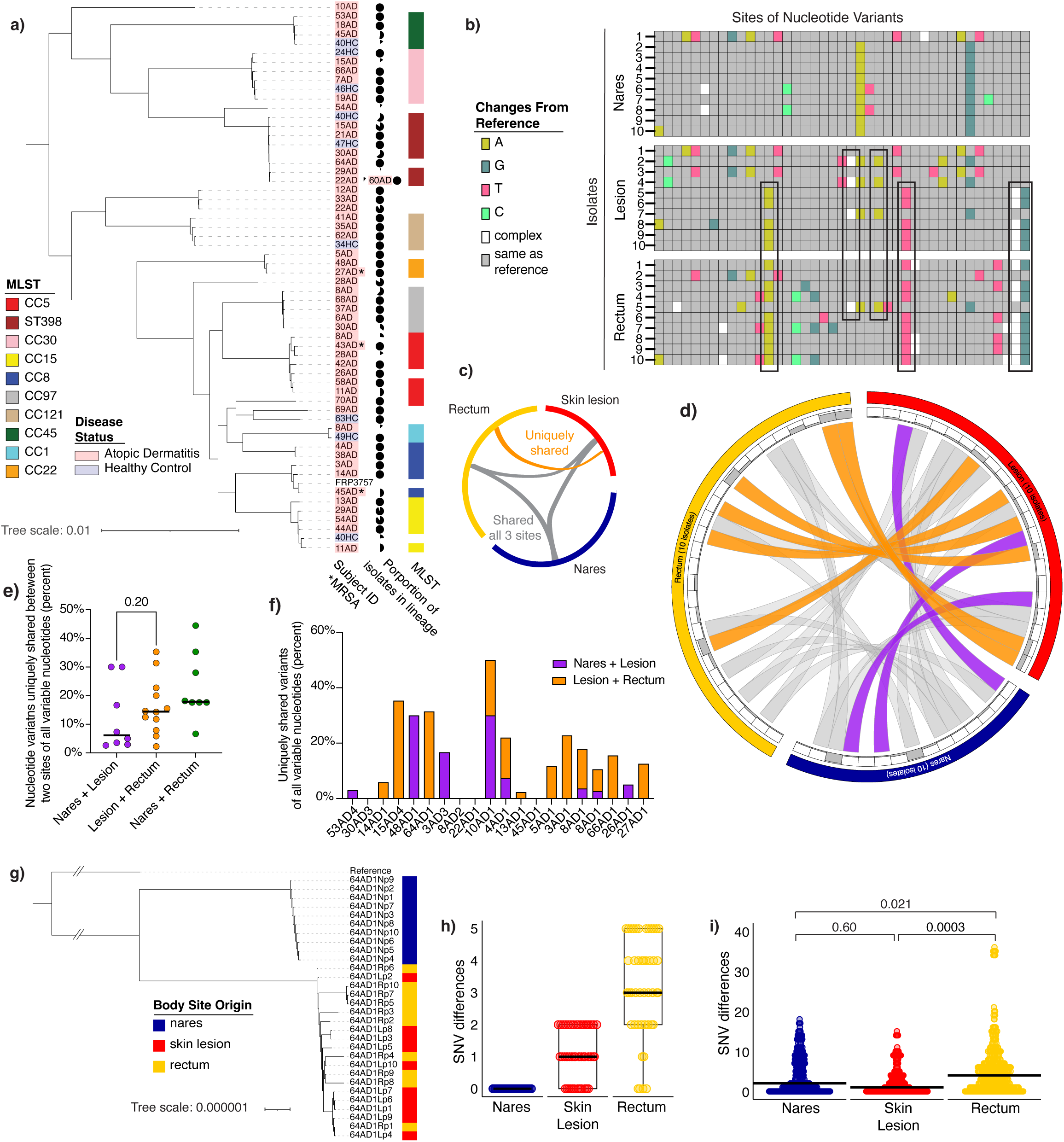
*S. aureus* is transmitted between the skin and gastrointestinal tract. **a)** Phylogenetic analysis of study strains from AD cohort. Maximum-likelihood phylogeny based on core genome of representative isolates from each lineage. Tips of the phylogenetic tree are colored according to the multi-locus sequencing type (MLST) and disease status and labelled with subject. Asterisks indicate methicillin resistant isolates (MRSA). Adjacent pie charts indicate proportion of isolates collected from each subject in that lineage. Note, subjects 22AD and 60AD cluster together using a 50 SNV threshold and, hence, are represented on the same branch; however, closer examination of the within-cluster SNV matrix, nevertheless, confirms that these two subjects have genomically distinct isolates (see **Methods** for further details). **b)** Heat map of sites of nucleotide variants (SNVs) for a representative subject (4AD). Each row represents an isolate, and each column represents a site of nucleotide variation. Sites consistent amongst all isolates are masked. Uniquely shared variants between isolates from the skin and rectal sites are highlighted with boxes. **c)** A model representation of the circular plot that follows, for reference. **d)** Shared nucleotide variants by body site. A circular plot for a representative subject (4AD), depicting uniquely shared variants across isolates from different sites. Outer arcs represent isolates from specific body sites. Inner arcs show the proportion of isolates with the variant at each nucleotide site (bar chart). Ribbons connect sites that share the variant: Gray ribbons show variants that are shared between all three sites, and colored ribbons show those uniquely shared between only two sites. **e)** Nucleotide variants shared between two different body sites as a percentage of all variable nucleotides within a lineage. Specifically, each point represents one lineage with at least 10 sequenced isolates and at least one from each body site. Black bars represent median. Statistical significance assessed by Mann-Whitney U test. **f)** Uniquely shared nucleotide variants by site pair at a single time point for a given individual. **g)** Maximum likelihood phylogeny of *S. aureus* isolates from one representative subject (64AD). **h)-i)** Within body site *S. aureus* genomic diversity. **h)** Pairwise differences in the number of SNVs between otherwise isogenic isolates from different body sites from a representative subject (64AD). Boxplot centered around median, lower and upper hinges indicating 1^st^ to 3^rd^ interquartile range (IQR), and whiskers indicating observations +/- 1.5 * IQR. **i)** Pairwise SNVs per isolate pairs across body sites for lineages with at least 7 isolates sequenced per site, summarized for all individuals (n = 10 subjects). Black line indicates median. Statistical significance assessed by permutation testing. SNV, single-nucleotide variant.

While previous studies have suggested a weak, geographically dependent correlation between certain *S. aureus* strains and AD, no specific sequence type has been consistently associated with the disease.^22,37^ In our study, we detected diverse *S. aureus* sequence types, none of which dominated in prevalence or were associated with disease severity (**Extended Data Fig. 4a-d**). These findings indicate that AD is not associated with any particular *S. aureus* clone and that most individuals carry unique lineages of the bacterium (**Fig. 4a**).

### The rectum is a reservoir for *S. aureus* in AD

To examine the within-host transmission dynamics of *S. aureus* among the nares, skin, and rectum, we focused on the 17 individuals who had at least one *S. aureus* isolate per site and a total of 10 or more isolates sequenced per timepoint. We assessed the unique and shared mutations across isolates from different body sites (example shown in **Fig. 4b**; additional cases in **Extended Data Fig. 5**). This analysis identified 63 instances of identical nucleotide variants exclusively shared between isolates from the lesional skin and rectal sites, compared to 31 such instances between lesional skin and nares sites. The shared variants in lesional skin and rectal isolates, detected in 11 of the 17 individuals analyzed, suggest direct *S. aureus* transmission between rectum and skin (examples in **Fig. 4c,d**; others in **Extended Data Fig. 6**).

To quantify these findings, we determined the proportion of variants shared exclusively between two body sites relative to all nucleotide variants observed within an individual at each timepoint. The median “exclusive sharing” rates were 6.2%, 14.5%, and 17.9% for nares-skin, rectum-skin, and nares-rectum pairs, respectively (**Fig. 4e**). No significant difference was observed between the frequency of variants shared exclusively by rectal-skin vs. nares-skin sites, indicating that the rectum contributes at least as much to *S. aureus* skin colonization as the nares. In 12 of the 17 individuals, variants were shared solely between rectum-skin or nares-skin pairs (**Fig. 4f**), suggesting that in these cases, cutaneous *S. aureus* originates from either the rectum or nares. Overall, these data support the hypothesis that *S. aureus* is directly transmitted between the rectum and skin, with transmission occurring as frequently from the rectum as from the nares.

Determining the direction of within-host *S. aureus* transmission is challenging, as many individuals were assessed at only a single time point. However, the distribution of isolates from the GI tract within individual phylogenies (example in **Fig. 4g**; others in **Extended Data Fig. 7**)— including intermingling with skin isolates and basal placement—suggests that, in some individuals, GI carriage represents a more established population compared to the other body sites and thus a potential source of *S. aureus* that colonizes affected skin. To further explore this possibility, we compared the genomic diversity of *S. aureus* within each body site to identify potential source sites. To ensure that enough isolates were available for comparison per individual, we focused on subjects having at least 7 isolates sequenced per site for all three body sites. Low or absent genomic diversity at a given site suggests recent seeding from another location, whereas sites with higher diversity likely serve as ancestral or reservoir sites.^37^ By counting nucleotide variant differences between *S. aureus* isolate pairs within each site, we found that in several individuals, the rectal site exhibited the highest genetic diversity compared to nares and lesional skin sites (see example in **Fig. 4h**, additional cases in **Extended Data Fig. 8**). Aggregated data across all subjects confirmed that the rectum harbored more genetically diverse *S. aureus* than the nares or lesional skin (*p=*0.021 and *p*=0.0003, respectively; **Fig. 4i**), suggesting that *S. aureus* has either diversified over a longer time within the rectum or that the rectum provides a more permissive environment for diversification. These findings indicate that the GI tract serves as a reservoir of *S. aureus* genetic diversity, acting as a potential source for transmission to the skin and contributing to AD pathogenesis.

### Adaptive mutations inactivate the *agr* quorum-sensing system

Examining both intra- and interhost genomic diversity can help identify host-driven adaptations and their potential role in AD.^22,34^ To explore this idea, we first analyzed our dataset for evidence of at least two mutations within the same gene across different isolates from the same individual, which would suggest within-host selective pressure. We focused on genomes from subjects with at least two sequenced isolates (*n*=913 isolates), regardless of body site. This analysis identified independent nonsynonymous mutations in 22 genes having known functions (**Fig. 5a**). In both AD and control isolates, mutations were noted in surface- and adhesion-related genes (e.g., *fbnB*, *sdrD, sdrE*) and in biofilm-formation genes (i.e., *hysA*). In contrast, unique mutations were enriched in the AD group in *agrC* (receptor histidine kinase of the *agr* quorum-sensing operon)^18^, *atl* (autolysin)^38^, *esaG1* (type 7 secretion system)^39^, and *rpsI* (ribosomal subunit). The occurrence of parallel mutations across isolates within individual patients (convergent evolution) suggests that *S. aureus* is adapting to the AD environment.

**Fig. 5:**
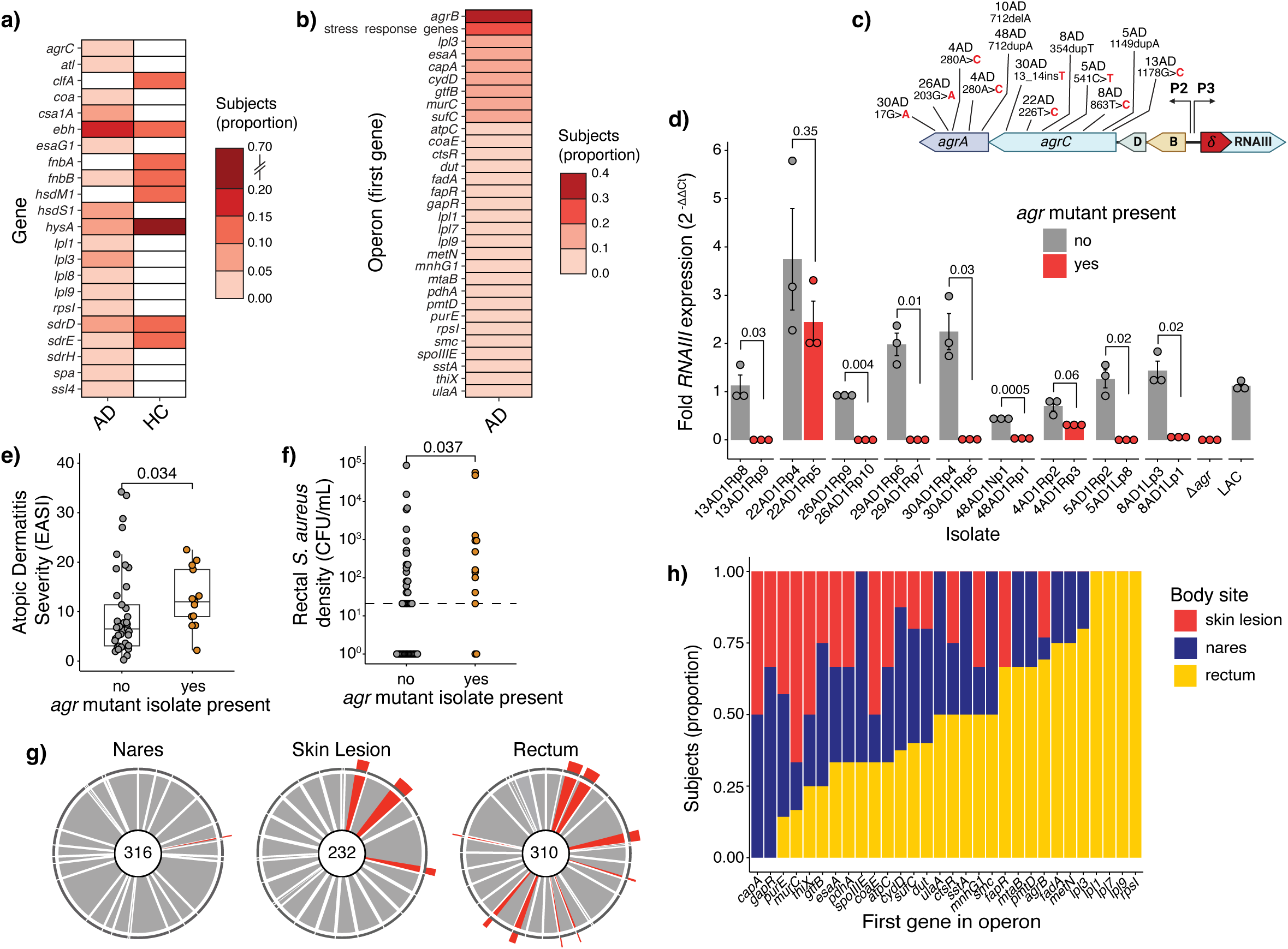
The gastrointestinal tract is site of *S. aureus* adaptive evolution in atopic dermatitis. **a)** Nucleotide variants by *S. aureus* gene. Proportion of subjects with nucleotide variants in each indicated gene by disease status (n = 37 AD, 7 HC subjects). **b)** Multiple nucleotide variants in *S. aureus* operons. Proportion of subjects with more than one nucleotide variant within the same operon. Operons observed in the HC group were excluded. Operons for which at least 3 independent subjects harbored mutations are shown (n = 37 AD subjects). **c)** Representative variation in the *agr* locus among 13 *agr*-defective strains. Shown are positions of nucleotide variants. **d)** *RNAIII* expression as functional output of *agr* locus. Normalized *RNAIII* expression in *agr*-WT vs. *agr*-mutated isolate pairs from the same lineage. LAC is shown as positive control and LACΔ*agr* as negative control. Significance tested by two-tailed t-test. **e-f)** AD severity and rectal *S. aureus* density by *agr* mutation status (*agrA* or *agrC*) at the time of sampling (n = 53; multiple timepoints included per subject, N = 37 AD subjects). **e)** AD severity measured by EASI in subjects with or without *agr* mutations **f)** *S. aureus* rectal density by *agr* mutation status. For **e)** and **f)**, statistical significance assessed by Mann-Whitney U test. **g)** *agr* mutation by body site. Pie charts per body site showing frequency of agr-WT (gray) or *agr*-mutant (red) isolates per study subject, as indicated by dark grey outer ring. Total number of isolates tested per site listed in center of pie. **h)** Distribution of mutated operons by site of origin. For each subject colonized by isolates with nucleotide variants from different body sites, the subject is represented once per body site. EASI, Eczema Area and Severity Index. AD, atopic dermatitis. HC, healthy control.

Next, we assessed variants both within and across AD subjects at the operon level, excluding the above-described surface protein- and biofilm-encoding genes that were also observed in controls. This approach identified mutations in the capsule locus, as previously reported^22^, validating our method (**Fig. 5b**). The most frequent mutations were putative inactivating mutations in the *agr* locus (12 of 36 subjects, **Fig. 5c**). To confirm attenuation of *agr* mutants, we demonstrated (1) reduced or absent production of δ-hemolysin (**Extended Data Fig. 9**), a toxin upregulated by *agr*^40^, and (2) decreased expression of *RNAIII*, the effector of the *agr* response (**Fig. 5d**).^18^

Our analysis also revealed mutations in operons associated with stress responses, such as the *clpC*^41^ and *sufCDSUB*^42^ operons. Given *clpC*’s role in managing misfolded proteins, we grouped the functionally related *clpB*, *dnaK/dnaJ,* and *groES/groL* operons^41,43^, along with *clpC*, as one regulon for analysis. Together, these stress response genes were the second most commonly mutated gene group (9 of 36 subjects, **Fig. 5b**). These findings suggest a pattern of adaptive mutations in pathways that regulate quorum sensing and certain stress responses that may contribute to *S. aureus* persistence in the AD environment. Supporting this idea, the presence of one or more *agr*-mutant *S. aureus* isolate was associated with more severe AD and increased rectal *S. aureus* density compared to subjects colonized exclusively by *agr*-sufficient *S. aureus* (*p*=0.034 and *p*=0.037 respectively, **Fig. 5e, f**).

Having identified adaptive mutations relevant to AD, we next evaluated the body-site origin of isolates carrying these mutations. Mutations in *agr* genes were most common in rectal isolates (**Fig. 5g**). In all cases where skin isolates harbored *agr* mutations, the rectum was also colonized with isogenic *agr* mutant *S. aureus*, but the reverse was not observed (**Extended Data Table 3**). These findings imply directionality, reinforcing the idea that the rectum can serve as a reservoir, seeding affected skin. Notably, *agr* mutants always appeared alongside wild-type *S. aureus* isolates (**Fig. 5g**). The exclusive presence of mixed populations within patients may reflect a recent allelic split. However, since this pattern was observed independently across multiple subjects, it suggests that the presence of wild-type organisms may facilitate colonization of the GI tract and skin by *agr* mutants in AD patients.

Broadening this analysis to mutated operons listed in **Fig. 5b** revealed that these mutations were similarly unevenly distributed by body site, further providing evidence of site-specific adaptation (**Fig. 5h**). Notably, despite comparable numbers of nares and rectal isolates analyzed (316 and 310 isolates respectively), the rectum was again overrepresented as a source or reservoir of adaptive mutations. Altogether, these data highlight body-site-specific adaptation in *S. aureus* and suggest that the GI tract plays a key role in emergence or maintenance of adaptive mutations, including those implicated in *S. aureus* pathogenesis in AD.

### Gastrointestinal-colonizing *S. aureus* increases skin inflammation in an infantile AD mouse model

Our human data suggest that GI colonization by *S. aureus* exacerbates skin pathology in AD. To directly test this hypothesis, we used a newly described infantile AD mouse model, in which dermatitis is induced by intradermal injection of house-dust mite (HDM; *manuscript currently under review*). We chose this model for its relevance to pediatric AD. Permissiveness of young mice to *S. aureus* GI colonization compared to adult mice^44^ parallels the human situation in which colonization resistance increases with age.^45^ Specifically, C57BL/6 pups were orally inoculated with *S. aureus* or phosphate-buffered saline (PBS), followed by intradermal HDM injection the next day to induce mild skin inflammation (**Fig. 6a**). We found that *S. aureus* GI tract colonization exacerbated skin inflammation, as evidenced by increased transepidermal water loss (TEWL, a measure of skin barrier integrity), compared to controls (*p*=0.004; **Fig. 6b,c**). Histologically, pups colonized by *S. aureus* demonstrated more severe signs of AD, including hyperkeratosis, acanthosis, and spongiosis, along with increased epidermal thickness (*p=*0.029, **Fig. 6e,f**). Additionally, transcriptional analysis of biopsies showed a 3-fold increase in *IL17a* expression in the *S. aureus-*treated group compared to controls, but no difference in Type 2 cytokine expression (**Fig. 6g**). These data align with the Th17-dominant immune response observed in early-onset pediatric AD^46^, and they suggest that *S. aureus* GI colonization contributes to Th17 skewing at distal skin sites. Collectively, these data support the hypothesis that *S. aureus* GI tract colonization exacerbates AD skin pathology in a causal manner.

**Fig. 6:**
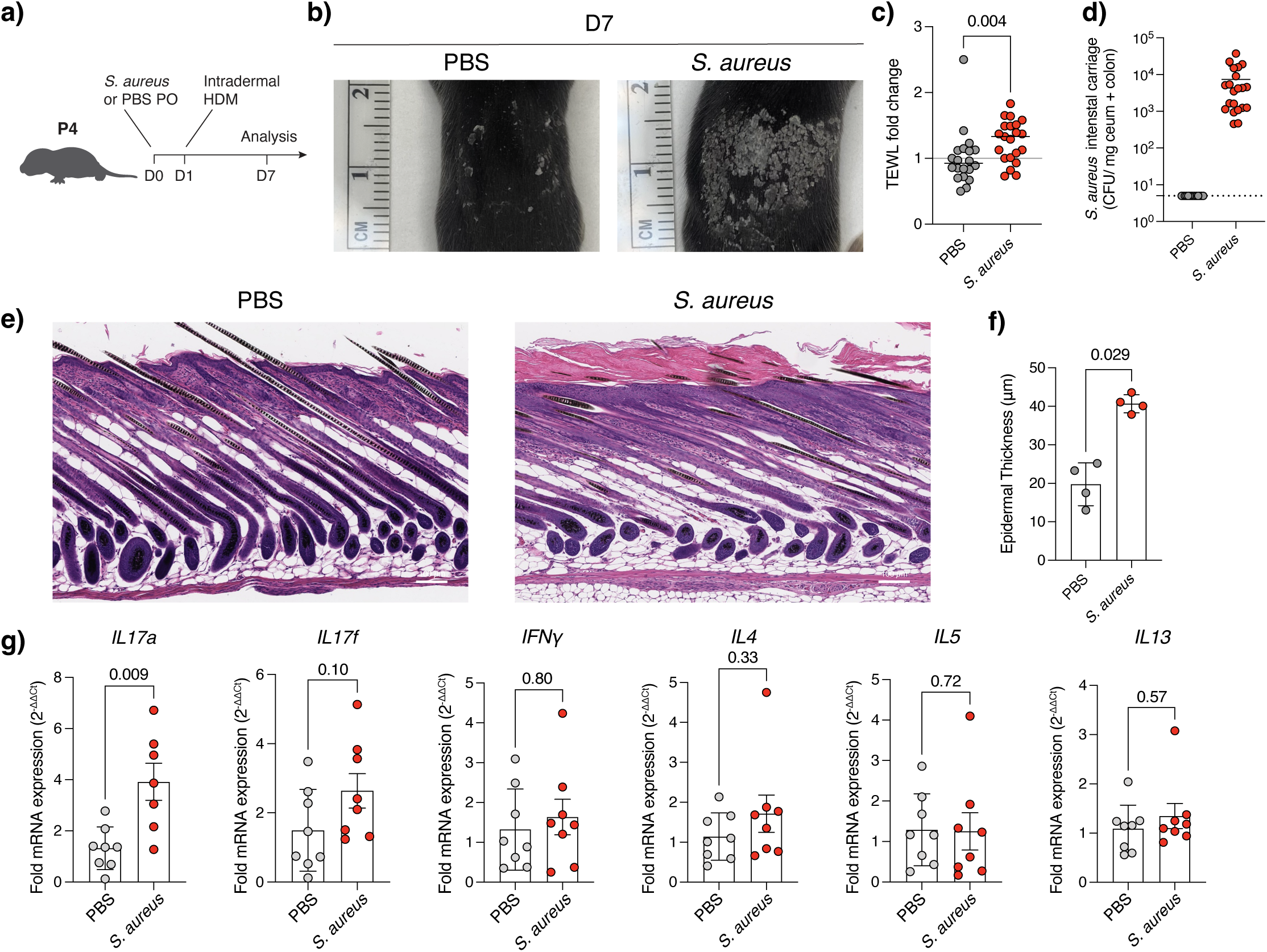
Gastrointestinal *S. aureus* exacerbates skin pathology in an infantile atopic dermatitis mouse model. **a)** Experimental design. Postnatal day 4 pups are gavaged orally with PBS or 1 x 10^7^ CFU *S. aureus* LAC (D0). The next day (D1), pups are intradermally injected with house-dust mite (HDM) to elicit a mild dermatitis. Six days following allergen injection (D7), pups are sacrificed for analysis. **b)** Representative images of dermatitis on D7 in pups orally inoculated with PBS vs. *S. aureus*. **c)** Transepidermal water loss (TEWL) on D7. Data represents three experiments (each with PBS and *S. aureus* group) performed on three separate days. TEWL expressed as fold-change relative to average TEWL of PBS group for that day (*n* = 20 PBS, 21 *S. aureus*). Black bars denote the median. Significance tested by Mann-Whitney U Test. **d)** *S. aureus* CFU count from harvested ceca and colon of pups on D7 (*n* = 20 PBS, 21 *S. aureus*). Dotted line at limit of detection of 5 CFU/mg cecum + colon. **e)** Representative histologic images of dermatitis on D7 in pups orally inoculated with PBS vs. *S. aureus*. Hemotoxylin-eosin stain. 10x magnification. Scale bar indicates 100 μM. **f)** Quantification of epidermal thickness from skin cross sections (*n* = 4 PBS, 4 *S. aureus*). Data represents one experiment. Significance tested by Mann-Whitney U test. **g)** Skin cytokine expression. Normalized cytokine expression of skin biopsies on D7 from pups orally inoculated with PBS vs. *S. aureus* (*IL17a*: n = 8 PBS, 7 *S. aureus*; *IL17f*: n = 8 PBS, 8 *S. aureus*, *IFNγ*: n = 8 PBS, 8 *S. aureus*, *IL4*: n = 8 PBS, 8 *S. aureus*; *IL5*: n = 8 PBS, 8 *S. aureus*, *IL13*: n = 8 PBS, 8 *S. aureus*). Data represents two independent experiments (each with PBS and *S. aureus* group) performed on two separate days. Significance tested by Mann-Whitney U test.

## DISCUSSION

In this study, combined sampling of the nares, skin, and rectum, along with validation in an independent cohort, revealed a previously unappreciated link between AD and *S. aureus* GI tract colonization. We found that rectal colonization by *S. aureus* is associated with AD presence and severity, highlighting the GI tract as a key colonization site in AD. Genomic analysis revealed that the GI tract serves as a reservoir of *S. aureus*, seeding affected skin and driving the selection of adapted *S. aureus* variants. Supporting these findings, mechanistic experiments in a murine model confirmed that GI-colonizing *S. aureus* increases AD skin pathology. Collectively, these insights indicate a novel AD-gut-*S. aureus* axis and could explain the limited efficacy of treatments targeting only cutaneous *S. aureus*.^47,48^

More broadly, our data contribute to the growing understanding that disruptions in local immune homeostasis at barrier sites lead to dysbiosis and pathogen enrichment.^49,50^ Recent attention has focused on the enrichment of *S. aureus* at skin sites due to microbiota alterations, such as depletion of skin-commensal coagulase-negative staphylococci.^21,51^ Our findings extend this concept beyond such local interactions, showing that microbial dysbiosis at an anatomically distinct site—the gut—can influence host-pathogen dynamics at a distal site, namely the skin, and exacerbate disease. Specifically, in AD subjects *S. aureus* GI colonization was associated with reduced gut microbiome diversity and the enrichment of *Corynebacterium* and other species. This dysbiosis may be a cause or consequence of the disease, as children with AD have previously been shown to have immature gut microbiomes.^29^ Dysbiosis likely facilitates the establishment of an *S. aureus* gut reservoir, which then seeds the skin, thereby exacerbating pathology. Upon AD resolution, *S. aureus* rectal density decreased, suggesting a reciprocal relationship between gut colonization and skin health, and highlighting a previously unexplored aspect of gut-skin immunomodulation.

Two key findings from our genomic analysis of *S. aureus* demonstrate that this gut-skin link is specific and likely directional in nature: We observe (1) exclusive sharing of SNVs between the gut- and skin-derived bacterial isolates – evidence for within-host transmission specifically between these two body sites; and (2) a greater diversity of *S. aureus* gut isolates compared to both the skin and nares. Higher genomic diversity in a particular niche suggests that that niche is more established and, therefore, ancestral to other niches or acting as a reservoir.^37^ A possible alternative explanation of this greater diversity among GI-colonizing isolates is that the gut niche is more readily captured than the skin due to its higher overall bacterial density. However, we restricted our analysis to individuals with at least seven *S. aureus* isolates per body site for comparison, making this less likely.

A closer examination of the mutations contributing to this *S. aureus* diversity revealed patterns of convergent evolution, suggesting that these mutations likely emerge as a product of the ecological pressures of the GI environment. Among these, loss-of-function mutations in the *agr* locus were the most common. These *agr* mutants, frequently found in the rectum and on the skin of AD patients, were rare in the nares, and they were associated with greater disease severity and higher bacterial density in the GI tract. In an apparent paradox, prior studies have linked *agr* activity to increased AD risk^20^ and found it to be required for GI colonization in animal models.^19^ However, a model of an “*agr* functionality trade-off” reconciles these findings: *agr* functionality may enhance initial GI colonization and AD development, but it may also disadvantage *S. aureus* under stress during AD flares. In this setting, *agr* inactivation may be required for persistence. For example, *agr* inactivation has been shown to provide a survival advantage in autophagic keratinocytes.^52^ Overall, these findings suggest the gut facilitates *S. aureus* strain evolution and, potentially, *S. aureus* adaptation to the fluctuating environment of waxing and waning AD.

In summary, our findings identify the gut as a relevant reservoir for *S. aureus* in AD and a dynamic niche that promotes *S. aureus* strain diversification. *S. aureus* GI colonization correlated with AD severity in humans, and it directly contributed to pathology in a murine AD model, underscoring its role in AD pathogenesis beyond that of an “innocent bystander.” Further, we observed that *S. aureus* GI colonization is associated with decreased gut microbiome diversity in children with AD and with an enrichment of specific co-colonizing species. Moving forward, establishing the mechanisms underlying this microbial gut-skin-axis and identifying *S. aureus*- and microbiome-specific therapeutic targets could pave the way for novel treatments for inflammatory skin conditions such as AD.

## METHODS

### AD study cohort and sample collection

Patients were recruited from the Pediatric Dermatology Clinic at the NYU Langone Ambulatory Care Center with approval by the Institutional Review Board (protocol no. i21-00426). Study subject demographics are provided in **Extended Data Table 1**. Inclusion criteria for enrollment in the AD arm were: 1) ages 6 months to 18 years, and 2) diagnosis of AD per modified Hannifin and Rajka Criteria,^53^ confirmed by a pediatric dermatologist. Inclusion criteria for enrollment in the healthy control (HC) arm were: 1) ages 6 months to 18 years, and 2) no known or suspected skin and soft tissue infections within the last 5 years and no history of AD. Exclusion criteria for both arms were: 1) pregnant subjects, children with impaired cognition, or HIV-positive individuals, 2) use of systemic or topical antibiotics within 3 months prior to enrollment, 3) fever > 38 °C, 4) use of immunosuppressive or immunomodulatory medications including (but not limited to) cyclosporine, methotrexate, mycophenolate mofetil, dupilumab, oral steroids within the last 30 days or 5 half-lives (whichever is longer), 5) phototherapy or light therapy within the last 30 days, 6) use of a bleach bath within the last 7 days, 7) other pruritic or inflammatory skin condition (e.g., scabies, ichthyoses, cutaneous T-cell lymphoma, psoriasis, photosensitivity dermatoses, immune deficiency diseases, or erythroderma of other causes). Written informed consent was obtained from parental guardians or participants aged 17 and older. Written informed assent using age-appropriate language was obtained from children aged 7-17 years.

For all subjects, bacteria were collected by swabbing the anterior nares, skin sites, and rectum (0.5 cm insertion for 15 seconds). For AD subjects, lesional swabs were collected from the antecubital fossa and unaffected skin from the abdomen. If no lesions were present in the antecubital fossa, samples were obtained from an involved area above the waist if present. Otherwise, any available skin lesion was sampled. AD disease severity was determined at time of sample collection by Eczema Area Severity Index (EASI: 0 = clear; 0.1-7.0 = almost clear/ mild; 7.1-21.0 = moderate; 21.1-72.0 = severe/ very severe).^54^ HC subjects had swabs taken from the antecubital fossa and abdomen.

Two swabs were collected from each site: one stored in culture media (1 mL of 3% tryptic soy broth [TSB] and 16.67% glycerol in phosphate-buffered *S. aureus* line [PBS]) to test for *S. aureus* growth, and the other in lysis buffer (1mL of molecular biology grade TE buffer [Invitrogen] containing 0.1% Triton X-100 and 0.05% Tween-20) for DNA extraction. For a random subset of subjects, a swab was waved in the air at the time of collection and subsequently stored in lysis buffer to assess for bacterial contamination. Swabs were stored at 4°C in solution for a maximum of 4 hours before transfer to the laboratory for storage at -80°C and further analysis.

Subjects received a mail-in kit (EasySampler® Stool Collection Kit, Cat. No. 127000A, GP Medical Devices, Denmark) to send a stool sample within two weeks of their visit. Stool samples were mailed at room temperature and, upon receipt, stored at -80°C for further analysis.

### CHES study cohort and sample collection

To validate our findings in a larger cohort, we obtained 352 stool samples in lysis buffer from infants aged 6 to 14 months enrolled in the NYU Children’s Health Environmental Study (CHES).^26^ Samples were collected using the OMNIgene®•GUT collection kit (Cat. No. OMR-200, DNA Genotek, Canada). Among these infants, 59 were diagnosed with AD based on a validated questionnaire^26^ (i.e., “Have you ever been told by your baby’s doctor that your baby has had eczema or dermatitis?”) administered at 12 months of age. The remaining 188 subjects were categorized as healthy controls.

### *S. aureus* culture

Swabs stored in TSB/glycerol were rapidly thawed, vortexed for 30 seconds, and a 50 μL aliquot of the solution was serially diluted, if necessary, and plated on selective CHROMagar**™** Staph aureus agar (Cat. No. TA672, Chromagar, France). Pink colony-forming units (CFU) were enumerated after overnight incubation at 37°C. From each swab, up to ten *S. aureus* colonies were isolated by subculture twice on tryptic soy agar (TSA) with 5% sheep blood and subsequently stored in TSB at -80°C for further analysis. For DNA extraction, cells were cultured on TSA plates.

### *Salinibacter ruber* spike-in preparation

*Salinibacter ruber* DSM 13855 (Cat. No. BAA-605, ATCC, Virginia) was spiked into samples for DNA extraction as previously described^30,31^ to obtain absolute bacterial counts from 16S rDNA gene sequencing. We cultured *S. ruber* by adding 50 μL of *S. ruber* stock to 25 mL of ATCC halobacterium medium 1270 in a sealed 250 mL Erlenmeyer flask, incubating for 24 days at 37°C. To further expand the culture, we transferred 4 mL of this culture to 250 mL of the same medium in a sealed 1-liter Erlenmeyer flask and incubated for 8 more days at 37°C. We concentrated cultures by centrifugation at 16,000 G for 15 min at 4°C, removed the supernatant, resuspended the pellet in PBS, and aliquoted it into 1 mL cryovials for storage at -80°C. Fresh aliquots were thawed, and 0.01 and 0.001 optical density units (ODU 600 nM) of *S. ruber* were added to each stool and rectal swab sample, respectively. *S. ruber* CFU per aliquot were enumerated by flow cytometry.

### Rectal swab and stool sample DNA extraction

DNA extraction from stool samples and rectal swabs was performed using the DNeasy PowerSoil Pro QIAcube Kit (Cat. No. 47126, QIAGEN, Maryland). Stool samples were aliquoted into pre-weighed homogenizer tubes and weighed. For rectal swabs, the swab head and 200 μL of the lysis solution were added to the homogenizer tubes. The appropriate amount of *S. ruber*, as described above, was added to the kit lysis buffer (CD1 solution) and mixed with each sample. For each DNA extraction, two negative controls were included: one without template material and one with *S. ruber* only (to assess purity of spike-in). DNA extraction was performed according to the manufacturers’ instructions and eluted in 100 μL of elution buffer.

### Droplet digital PCR (ddPCR)

We designed a ddPCR assay directed against the *S. aureus*-specific *spa* gene using previously published primers^55^ and a custom probe (see **Extended Data Table 4**). To assess sensitivity, we evaluated the primer binding *in silico* using the amplicon command of seqkit^56^ v. 2.3.0 against our laboratory’s library of sequenced *S. aureus* genomes and confirmed effective binding (>99% with perfect matches to primers and probe). The specificity of the of the primers was validated using the NCBI Primer Blast tool targeting the RefSeq Representative Genome Database (Organism: Bacteria).^57^ We performed *spa*-ddPCR on rectal swabs from which *S. aureus* was cultured, confirming a correlation between the two (Pearson correlation, *R* = 0.8, *p* < 0.0001, **Extended Data Fig. 1b**). *spa*-ddPCR was performed in triplicate using the QX200 ddPCR system (Bio-Rad, California) and involved four steps: 1) preparation of the reaction mixture using ddPCR Supermix for Probes (No dUTP, Cat. No. 1863024, Bio-Rad, California) and AvrII restriction enzyme (Cat. No. R0174L, New England Biolabs, Ipswich, MA) with 2.4 μL of sample and final reaction volume 24 μL; 2) droplet generation using the Automated Droplet Generator (Bio-Rad, California), 3) ddPCR amplification with a cycling protocol of 95°C for 10 min, 40 cycles of 94°C for 30 sec and 60°C for 1 min, and 98°C for 10 min for enzyme deactivation; 4) droplet read and analysis on QuantaSoft 1.7.4.0917 software. Samples with less than 1 DNA copy/ μL detected were zeroed based on DNA extracted from serial dilutions of LAC culture demonstrating detection is lost at 1 DNA copy/ μL equivalent to 1 CFU/ μL. DNA copies obtained were then normalized to input amount of stool.

### 16S rDNA gene sequencing

Automated amplicon library preparation was performed on a Biomek 4000 using a custom liquid handling protocol. The V4 region of the 16S rDNA gene was amplified using barcoded primers.^58^ Each unique barcoded amplicon was generated in pairs of 25 μL reactions with the following reaction conditions: 8.5 μL PCR-grade H_2_O, 12.5 μL GoTaq G2 Master Mix (Cat. No. M7822, Promega, Wisconsin), 2 μL each of barcoded forward and universal reverse primers (5 μM) and 2 μL 100x diluted template DNA. Reactions were run on a C1000 Touch Thermal Cycler (Bio-Rad, California) with the following cycling conditions: initial denaturing at 94°C for 3 min followed by 35 cycles of denaturation at 94°C for 45 sec, annealing at 50°C for 1 min, and extension at 72 C for 90 sec, with a final extension of 10 min at 72°C. Amplicons were quantified using the 2200 TapeStation system (Agilent, California) and Quant-iT 1x dsDNA HS Assay Kit (Cat. No. Q3323, Invitrogen, Massachusetts) and pooled equimolar. Purification of the pool was then performed using 0.8x AMPureXP beads (Cat. No. A63880, Beckman Coulter, California) as per the manufacturer instructions and final concentration determination was determined using Qubit (Invitrogen, Massachusetts). Sequencing was performed using paired-end 150 cycles on a MiSeq (Illumina) using custom spike-in Read 1/Index/Read 2 primers.

### 16S rDNA bioinformatic processing and taxonomic assignment

Amplicon sequence variants (ASVs) were generated via QIIME 2^59^ v. 2023.5 using post-QC FASTQ files. Within the package, DADA2^60^ v. 2024.5.0 was used to denoise FASTQ reads trimmed to 150 base pairs. For all samples, a minimum of 10,000 denoised and non-chimeric reads were achieved and, therefore, all samples were included in further analysis. ASV taxonomy was assigned using the QIIME 2 feature-classifier^61^ v. 2024.5.0 and SILVA^62^ v.138 database. Reads assigned to *mitochondria* or *chloroplast* were excluded from further analysis. Downstream preprocessing was conducted in R using the phyloseq^63^ v. 1.48.0 package.

### Microbiome diversity and composition

*α-diversity. Salinibacter ruber* reads were excluded from this analysis. Using the phyloseq package, we calculated the inverse Simpson index from raw ASV reads.

*Principal component analyses. Salinibacter ruber* reads were excluded from this analysis. Using the phyloseq package, we agglomerated reads at the genus level and transformed the reads to relative amounts per sample. These reads were then used to calculate Bray-Curtis distances. The principal components of the resulting matrix were calculated, and sample compositions were plotted on the first two principal components. PERMANOVA was performed using the vegan^64^ v. 2.6.8 package.

*Microbial differential abundance analysis*. The absolute abundances of bacterial taxa were determined using previously described methods with minor modifications.^30,31^ In brief, the density of the *Salinibacter ruber* culture (spike-in bacterial species) was determined by flow cytometry. 1.05 × 10^5^ *S. ruber* cells were added to each rectal swab and 1.05 × 10^6^ cells to each stool sample prior to DNA extraction (see above). Raw counts were transformed into relative abundance. A pseudocount equivalent to half the minimum non-zero value across all samples was added to each sample. The absolute abundance of a given taxon *i* in sample *j* was then calculated as follows:

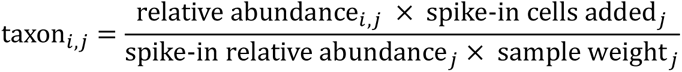

*S. ruber* reads were then removed from all samples. Taxa with less than 10% prevalence across all samples in a cohort were excluded. To test for associations of particular taxa with *S. aureus* density, we utilized the following linear mixed-effects model using the R package lmerTest^65^ v. 3.1-3 for a given a given taxon *i* in sample *j*. Note for several subjects, multiple samples were available from multiple timepoints.

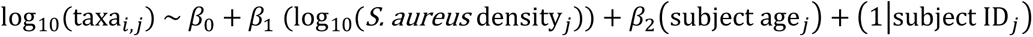

### *S. aureus* DNA extraction, whole genome sequencing, and bioinformatic processin

*Sequencing and Quality Control*. DNA was extracted from each *S. aureus* isolate as previously described.^66^ Briefly, DNA was extracted using the KingFisher Flex Purification System (Thermo Fisher, Massachusetts) quantified with the Quant-it picogreen dsDNA assay kit (Cat. No. P7589, Invitrogen, Massachusetts), and normalized by concentration. Libraries were prepared using the Illumina DNA prep (M) Tagmentation kit (Cat No. 20018705, California), pooled equimolarly, and sequenced as paired-end 150-bp reads using the Illumina Novaseq 6000 System with the S1 300 cartridge and flow cell (Illumina, California). The resulting paired-end 150 bp reads were filtered and trimmed using fastp^67^ v. 0.20.1. For quality control, the software ConFindr^68^ v. 0.7.4 was used to screen for within-species (strain mixture) contamination (exclusion threshold: >10% contamination) and MetaPhlAn^69^ v. 3.0.13 was used to screen for cross-species contamination (exclusion threshold: >5% contamination). The cleaned reads output by fastp were assembled using Unicycler^70^ 0.4.8 in conservative mode. The resulting assemblies were further examined using GTDB-Tk^71^ v. 1.5.1 to ensure that they were classified as *S. aureus*. The MLST software was used to determine *S. aureus* sequence types.^72^

*Assignment of S. aureus to lineages*. Filtered reads from the 1,157 isolates passing quality control were mapped to a genome assembly of *S. aureus* strain FPR3757 (NCBI RefSeq accession number GCF_000013465.1) using snippy^73^ v. 4.6.0; these assemblies were then combined with the snippy-core command, yielding a 2.13 Mbp core genome alignment. The snp-dists command of snippy was then used to generate a SNV distance matrix. The igraph^74^ v. 2.1.2 R package was used to generate an undirected graph in which vertices represent isolates and edges link two nodes whenever isolates differ by fewer than than 50 SNVs. The 50 SNV cutoff was empirically selected after observing that nearly all isolates differing by fewer SNVs were derived from the same individual (**Extended Data Fig. 10a,b**). Only isolates from subjects 22AD and 60AD clustered together below the 50 SNV threshold; however, closer examination of the within-cluster SNV matrix confirms that these two subjects have genomically distinct isolates (**Extended Data Fig. 10c**). Lineages were defined as the connected components of this network.

*AD cohort phylogenetic tree.* snippy was used to create a core genome alignment including one representative isolate from each lineage, using the FPR3757 reference genome. *IQ-TREE*^75^ v. 2.3.6 was then used to generate a phylogenetic tree using the GTR+F+G4 model of sequence evolution. To label isolates as methicillin-resistant, presence of the *mecA* gene as well as *mecA* type were determined using staphopia-sccmec v. 1.0.0.^76^

*Within lineage S. aureus core alignment*. To resolve genetic differences within isolate lineages, we again used snippy to generate a new core alignment for each cluster, using as reference assembly the RefSeq genome assembly with the highest average nucleotide identity (ANI) to the isolates in each cluster as calculated by FastANI v. 1.32.^77^

*Within lineage S. aureus genomic diversity analysis.* Distance matrices consisting of within-cluster SNV counts were used to assess differences in genomic diversity between body sites. Specifically, the median number of SNVs was calculated over all pairs of isolates from the same body site, individual, lineage, and timepoint. To compare the medians between two body sites, permutation testing was performed: 10,000 null datasets were generated in which site labels were randomly reassigned to isolates. For each comparison, the *p*-value was calculated as the fraction of null datasets in which the difference in medians exceeded that observed in the true dataset.

*Across subject S. aureus core alignment*. To facilitate comparison of mutated genes across subjects, we also used a single reference genome assembly (strain FPR3757 with RefSeq accession number GCF_000013465.1) to call variants within each cluster. Specifically, fastp-cleaned reads were aligned to the reference using bwa-mem v. 0.7.18.^78^ Joint variant calling was then performed on the aligned reads of all isolates belonging to each lineage using FreeBayes v. 1.3.2^79^ with options `-p 1 --min-alternate-count 10 --use-best-n-alleles 4 -- min-alternate-fraction 0.2 --genotype-qualities`. The resulting joint VCF file was filtered using bcftools v. 1.15.1^80^ keeping only sites that had genotypes called with genotype quality at least 100 for every isolate in a lineage. VCF files were further filtered to keep only sites with more than one allele called amongst the isolates in a cluster, since we were only interested in mutations that arose within a subject’s population of bacteria. This resulted in a joint VCF file for each lineage of isolates. These joint VCF files were then annotated using *snpEff*^81^, which was run with options `-noLog -noStats -no-downstream -no-upstream -no-utr` using a custom database generated for the strain FPR3757 assembly.

### *S. aureus* RNA extraction

*S. aureus* isolates stored in TSB at -80°C were streaked onto TSA with 5% sheep blood and incubated overnight at 37°C. To 400 μL of TSB in a deep well plate (Cat No. 1896-2110, USA Scientific Inc, Florida), 3 colonies of each isolate were added to each well in triplicate. The plate was sealed (Cat. No. 4306311, Applied Biosystems, Massachusetts) and incubated overnight shaking at 37°C. Subculture was performed on 5 μL of the overnight culture added to 400 μL of TSB in a fresh deep-well plate, sealed and incubated for 4.5 hours shaking at 37°C. The MagMAX™-96 Total RNA Isolation Kit (Cat. No. AM1830, Thermo Fisher, Massachusetts) was used with the KingFisher Flex Purification System (Thermo Fisher, Massachusetts) according to the manufacturer’s instructions with the following modifications: the subculture was spun down at room temp. for 10 min. at 4,000 G to form a bacterial pellet; supernatant was removed and 0.1 mm silica beads (Cat. No. 11079110z, BioSpec Products, Oklahoma) and 154 μL of Kit Lysis Binding solution were added and homogenized; the supernatant was transferred to a fresh deep-well plate and the protocol was continued as per the manufacturer’s instructions.

### *RNAIII* RT-qPCR

Complementary DNA was synthesized according to the manufacturer’s instructions (Maxima First Strand cDNA Synthesis Kit, Cat. No. K1641, Thermo Fisher, Massachusetts). Quantitative polymerase chain reaction (qPCR) was performed using SYBR Green PCR master mix (Cat No. 208054, QIAGEN, Maryland). Previously published primers were used for *RNAIII*^82^ and the housekeeping gene *rpoB*^83^ (see **Extended Data Table 4**).Three independent biological samples of each strain were run in duplicate and *rpoB* was used to normalize gene expression. Settings on the C1000 CFX96 machine (Bio-Rad Laboratories) were as follows: 50°C for two min, 95°C for 10 min, then 40 cycles [95°C for 15s and 60°C for 1 min]. The 2^-ΔΔCT^ method was used to calculate the relative fold gene expression.^84^ LAC^85^ and LACΔ*agr*^86^ were used as positive and negative control stains respectively.

### Early life atopic dermatitis mouse model

All experiments involving live animals were performed with approval of the Institutional Animal Care and Use Committee of NYU Grossman School of Medicine (protocol no. IA16-00087) and on C57BL/6J mice (Strain #: 000664, The Jackson Laboratory, Maine). Mice were housed in specific pathogen-free conditions with a 12-h light:12-h dark cycle at 20–22°C and 30–70% humidity. To evaluate the effect of GI-colonizing *S. aureus* on early life AD, we utilized a murine infantile AD model. Methods were provided by author correspondence as the manuscript describing the new model is still under review. Since day-to-day variability in humidity and temperature can affect transepidermal water loss (TEWL) measurements (see below), all murine experiments were performed on a minimum of 2 litters born on the same day to ensure an adequate number of pups for experimental and control groups treated in parallel. To avoid cage-effects, the litters were distributed across a minimum of 4 cages with birth and foster dams and with at least 2 experimental cages and 2 control cages per experiment.

### Transepidermal water loss (TEWL) measurement

On P11 (6 days following HDM treatment), mice were euthanized, the skin dissected and spread epidermal side up for TEWL measurement using the Tewameter ® Hex probe (Courage + Khazaka electronic, Germany). Data were collected for 5 minutes without disturbance. The TEWL values were calculated as the average TEWL robust (g/m^2^/h) value per mouse between 3’30’’ and 4’ of measurement, when the reading had stabilized, and normalized to the control cohort (PBS-treated) average measured on the same day.

### Murine histology

Mouse full-thickness skin biopsies were fixed in 4% paraformaldehyde overnight at 4°C, washed three times in PBS, and transferred to 70% ethanol for processing, embedding, sectioning, staining with hematoxylin and eosin, and imaging.

### Murine skin mRNA measurement

Mouse full-thickness skin biopsies were flash frozen in liquid nitrogen and stored in -80°C until RNA extraction. Total RNA was extracted using the RNeasy Mini Kit (Cat. No. 74104, QIAGEN, Maryland). Complementary DNA was synthesized according to the manufacturer’s instructions (Maxima First Strand cDNA Synthesis Kit, Cat. No. K1641, Thermo Fisher, Massachusetts). Quantitative polymerase chain reaction (qPCR) was performed in triplicate using SYBR Green PCR master mix (Cat No. A25778, Thermo Fisher, Massachusetts). Settings on the QuantStudio™ 6 Pro machine were as follows: 50°C for two min, 95°C for 10 min, then 40 cycles [95°C for 15s and 60°C for 1 min]. Analytes were normalized to housekeeping gene, *ACTB*. 2^-ΔΔCT^ method was used to calculate the relative fold gene expression.^84^ For complete list of qPCR primers, refer to **Extended Data Table 4**.

### Quantification and statistical analyses

Group sizes were determined based on results of preliminary experiments. Statistical significance was determined as shown in each figure legend. Statistical analysis was performed with R v. 4.3 and GraphPad Prism v. 10.4.0.

### Data Availability

All data to support the conclusions in this manuscript can be found in the main text, extended data, and supplemental materials. Sequencing data will be deposited in GEO. Manuscript will be updated with accession codes when available. Raw data from our study may be requested from the corresponding authors.

### Code Availability

All data were generated using previously published packages. Upon request, computer code will be made available to readers.

## Author Contributions

TKK and BS conceptualized the study. TKK, BS, VJT, AG, LT, and SN designed experiments. TKK, JShenderovich, CDN, AT, and VO recruited human study subjects. TKK, JShenderovich, AT, and CN enrolled human study subjects and executed clinical protocols. TKK, JShenderovich, SMM, AS, NA, AT processed samples. TKK, GP, and AP analyzed experimental data. JSchluter designed the microbiome analysis. TKK, GP, NMS and AP performed bioinformatic analysis. TKK, MP, AA, YX performed experiments. TKK and BS wrote the initial manuscript draft. All authors contributed during manuscript revision and approved the final version.

## Acknowledgements

We thank all study participants who devoted time to our research; Lorna Thorpe, Sarah Conderino and all Shopsin lab members for helpful discussions; Seth Rakoff-Nahoum and lab members for guidance on implementation of absolute 16S rDNA sequencing approach; David Polsky and lab members for guidance on ddPCR; NYU CHES study participants, and NYU CHES study staff; Karl Drlica for his reading of the manuscript. This work was supported by the National Institute of Allergy and Infectious Diseases at the NIH 5R01AI140754 and 5R01AI137336 (BS and VJT), K08AI163457 (RJU); the National Institute of Arthritis and Musculoskeletal and Skin Diseases at the NIH T32AR064184 (TKK), K08AR084045 (TKK); the Environmental influences on Child Health Outcomes (ECHO) Program, Office of the Director, National Institutes of Health, under Award Number UG3/UH3OD023305 (LT); the Pediatric Dermatology Research Alliance Travel Scholarship Award (TKK); the National Eczema Association Engagement Research Grant (TKK); the American Skin Association Leo Pharma Research Grant in Atopic Dermatitis (TKK); the Rudin Family Pilot Grant Program for Resident Research (TKK); the Childhood Eczema Prevention Research Grant (TKK); and funds from the NYU Langone Health Antimicrobial-Resistant Pathogens Program (BS, AP, and VJT). JSchluter reports funding from the NIH grants DP2 AI164318-01, and R01CA269617. SN reports funding from NIH grants DP2AR079173-01 and R01-AI168462, Burrow Welcome Path Award (24-2108) and the Leo Foundation (LF-AW_RAM-24-400348). S.N. is a NYSCF Robertson Stem Cell Investigator, a Packard Fellow, and an Allen Discovery Center for Neuroimmune Interactions Investigator. The content is solely the responsibility of the authors and does not necessarily represent the official views of the National Institutes of Health. The authors would like to acknowledge the NYU Langone Health Genome Technology Center (RRID:SCR_017929). This publication made use of the PubMLST website (https://pubmlst.org/) developed by Keith Jolley <u>(Jolley & Maiden 2010, BMC Bioinformatics, 11:595)</u> and sited at the University of Oxford. The development of that website was funded by the Wellcome Trust.

## Competing interests

SN is on the scientific advisory board of Seed Inc., cofounder of Stara Biosciences, and receives funding from Takeda Pharmaceuticals. JSchluter is co-founder of Postbiotics Plus Research and holds equity, serves on an advisory board and holds equity of Jona Health, and has filed intellectual property applications related to the microbiome (reference numbers # PCT/US2023/060616). VJT has received honoraria from Pfizer and MedImmune and is an inventor on patents and patent applications filed by New York University, which are currently under commercial license to Janssen Biotech Inc. Janssen Biotech Inc provided research funding and other payments associated with a licensing agreement not related to this study. TKK, GP, MP, JShenderovich, AT, SMM, AG, AS, NA, HS, AA, YX, NMS, CDN, LT, VO, AP, RJU, and BS report no relevant conflicts of interest.

---

**Extended Data Fig. 1:**
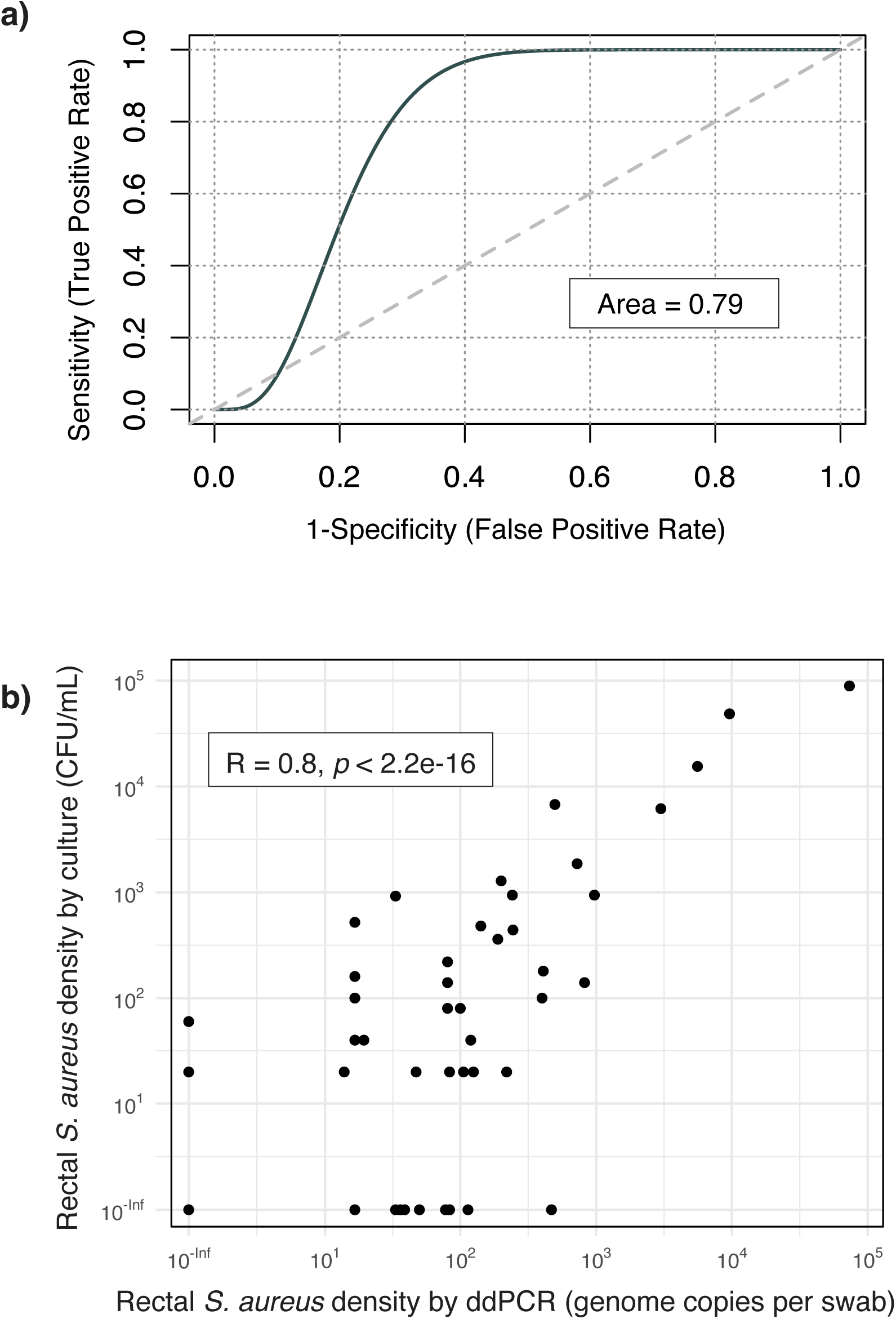
Comparison of *S. aureus* rectal swab culture with ddPCR from matched stool and rectal swabs. **a)** Receiver operating characteristic (ROC) curve comparing *S. aureus* density as detected by culture of rectal swab (detected vs. not detected) vs. ddPCR for *S. aureus*-specific *spa* gene from matched stool samples (genome copies per 100 mg stool); n = 27 paired samples). ROC curve plotted using binomial model. Area under curve = 0.79. Model applied and plot generated using ROCit package version 2.1.2.^87^ **b)** Correlation of *S. aureus* rectal density by ddPCR vs. culture from matched swabs (n = 88 paired samples). Significance assessed by Pearson correlation.

**Extended Data Fig. 2:**
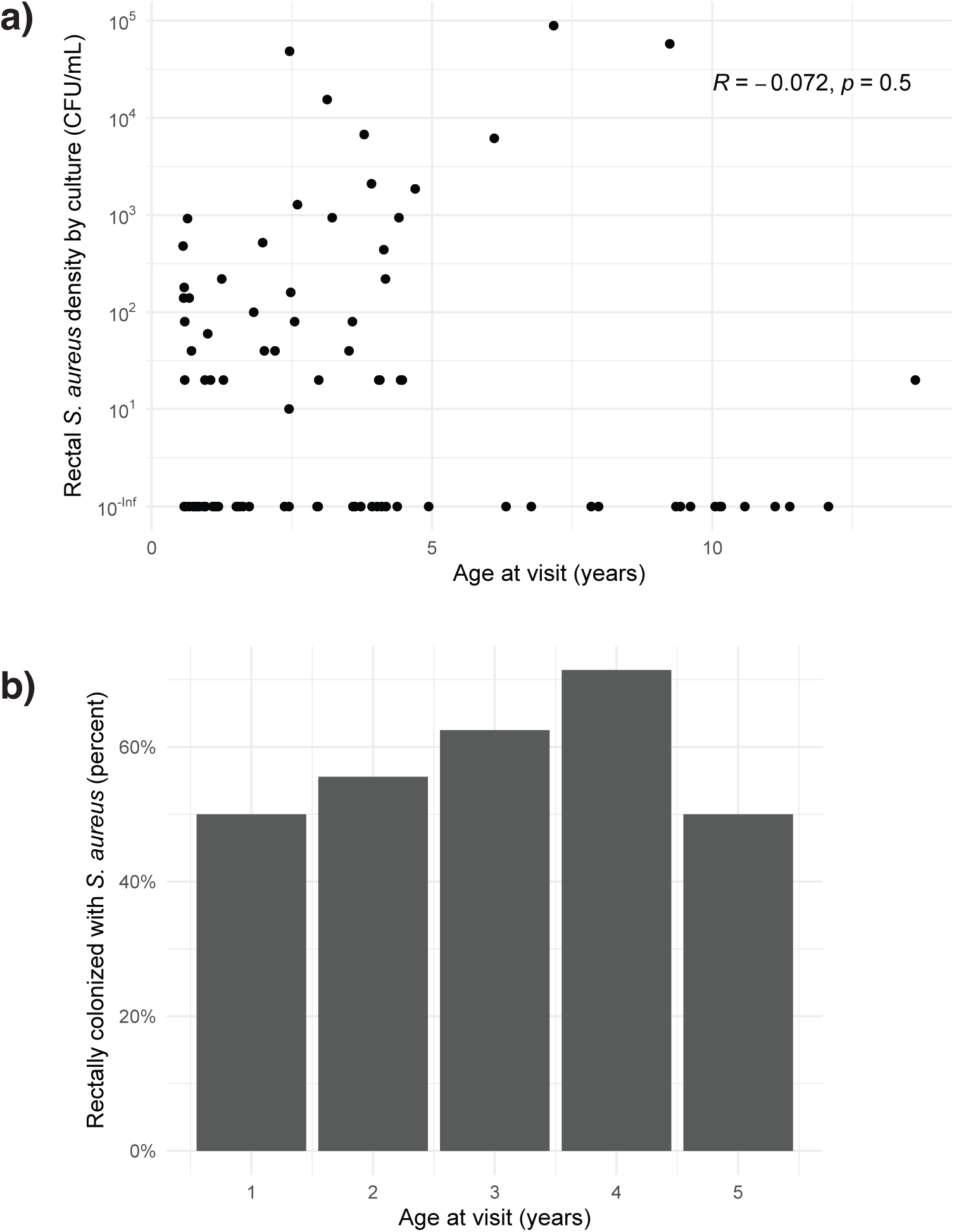
*S. aureus* rectal colonization by age in AD subjects. **a)** AD cohort: age at visit vs. rectal *S. aureus* density by culture; significance assessed by Pearson correlation (n = 93 samples, N = 61 AD subjects). **b)** AD cohort: proportion of subjects colonized rectally with *S. aureus* by culture binned by age groups (group = 1 year; N = 46 AD subjects; only first timepoint data included). Subjects older than 6 years old not included as there were less than four subjects per age group older than 6 years.

**Extended Data Fig. 3:**
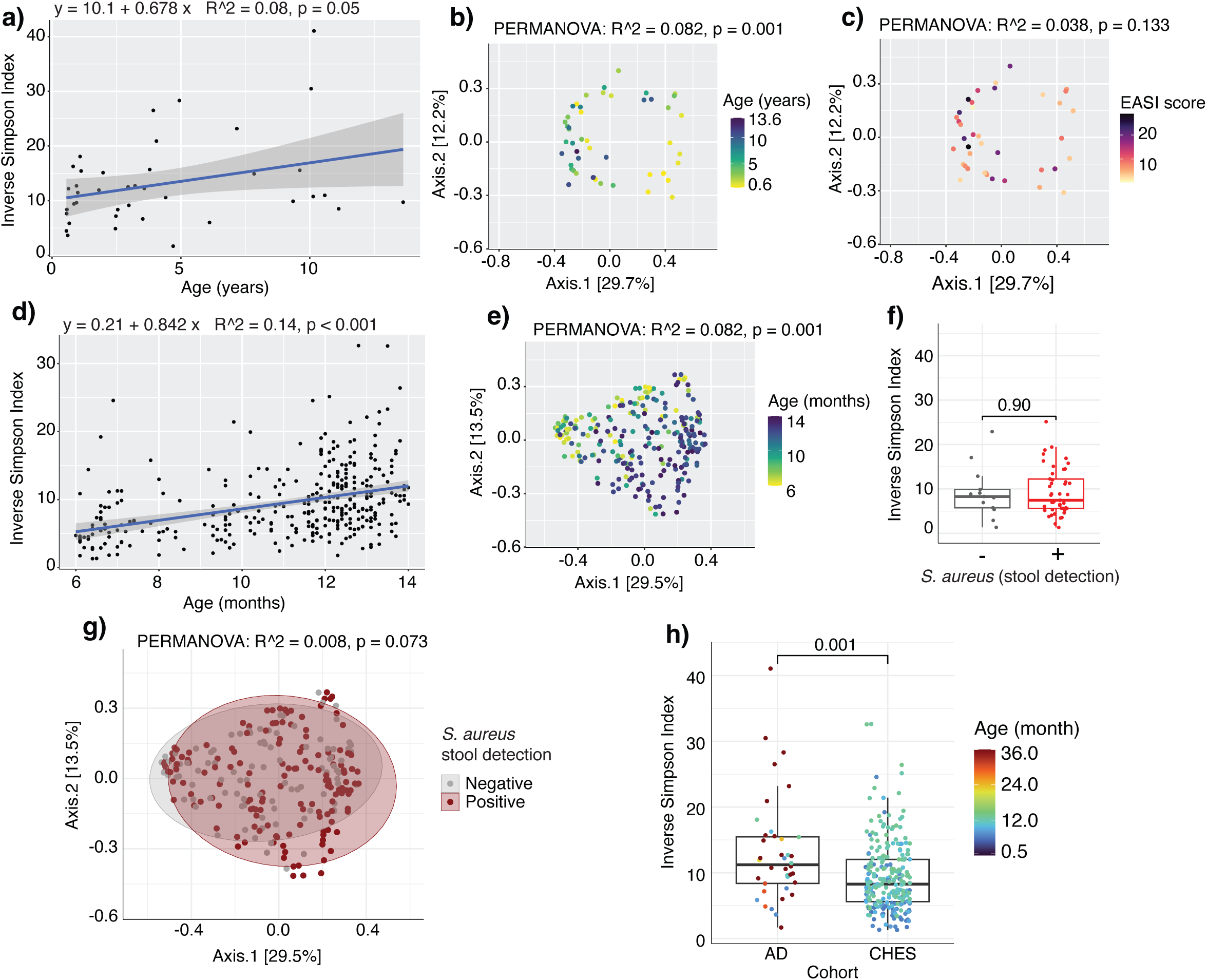
Effect of age and other parameters on gut microbiome. AD cohort (n = 40 AD, 9 HC subjects) **a)-c). a)** Linear regression of subject age and inverse Simpson index (R^2^=0.08, *p*=0.05). PCoA of Bray-Curtis dissimilarity by **b)** age (PERMANOVA, *p*=0.001) and **c)** AD severity as assessed by EASI score (PERMANOVA, *p*=0.133). CHES cohort (n = 58 AD, 188 HC subjects) **d)-g).** Linear regression of subject age and inverse Simpson index (R^2^=0.14, *p*<0.001). **e)** PCoA of Bray-Curtis dissimilarity by age (PERMANOVA, *p*=0.001). **f)** Inverse Simpson index by *S. aureus* stool detection by ddPCR (Mann-Whitney U test, *p*=0.90). **g)** PCoA of Bray-Curtis dissimilarity by *S. aureus* stool detection by ddPCR (PERMANOVA, *p*=0.073). **h)** Comparison of alpha diversity by inverse Simpson index of subjects from AD vs. CHES cohorts. Dots are colored by age of subject (Mann-Whitney U test, *p*=0.001). Inverse Simpson index assessed at ASV level. Bray Curtis dissimilarity calculated at genus level. AD, atopic dermatitis. EASI, Eczema Area and Severity Index.

**Extended Data Fig. 4:**
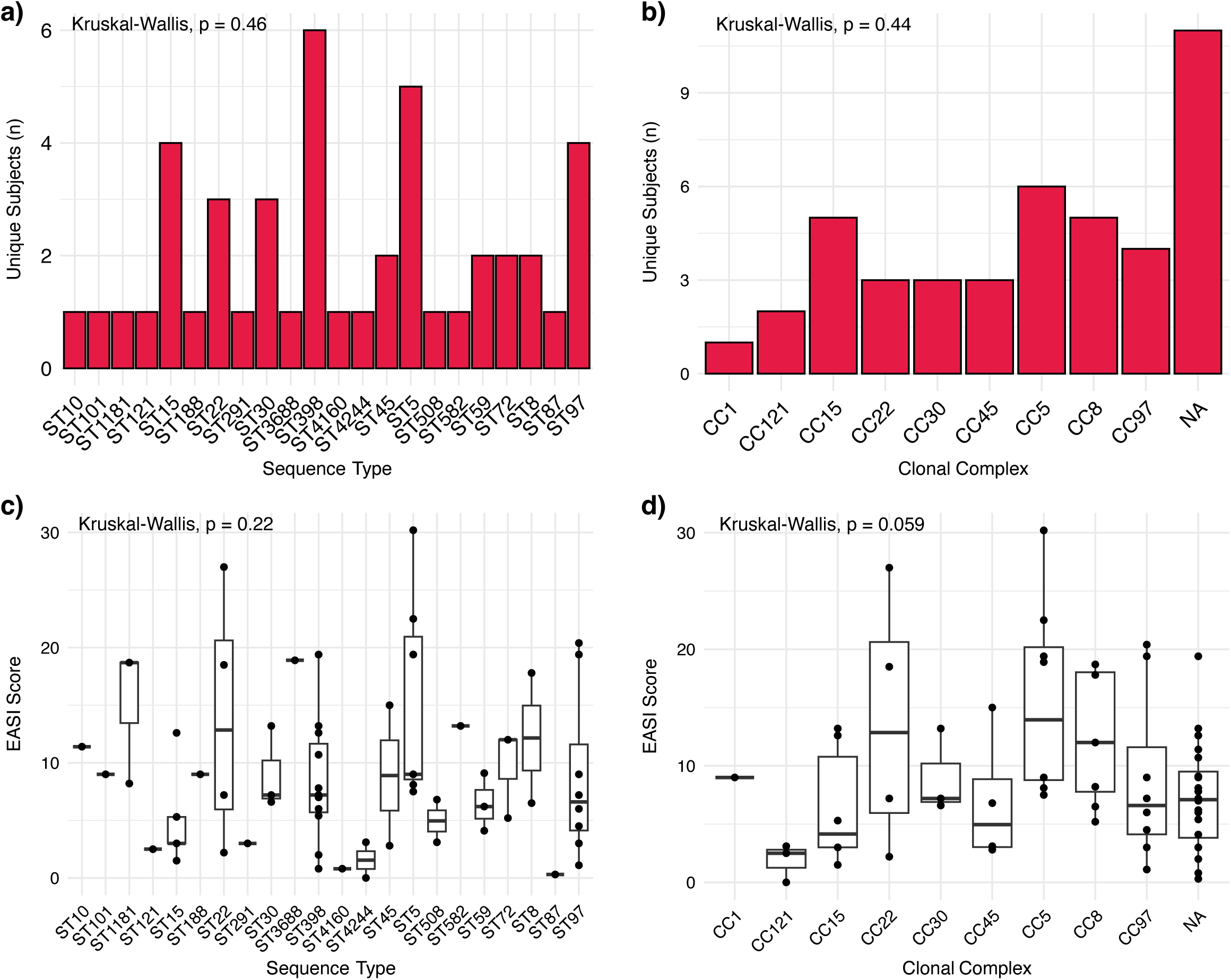
*S. aureus* sequence types by number of subjects and disease severity. *S. aureus* sequence types **a)**, **c)** and clonal complexes **b)**, **d)** colonizing AD subjects from AD cohort by number of subjects **a)**, **b)** and disease severity **c)**, **d)** as assessed by EASI score. Significance assessed by Krukal-Wallis test. Boxplot centered around median, lower and upper hinges indicating 1^st^ to 3^rd^ interquartile range (IQR), and whiskers indicating observations +/- 1.5 * IQR. AD, atopic dermatitis. EASI, Eczema Area and Severity Index.

**Extended Data Fig. 5:**
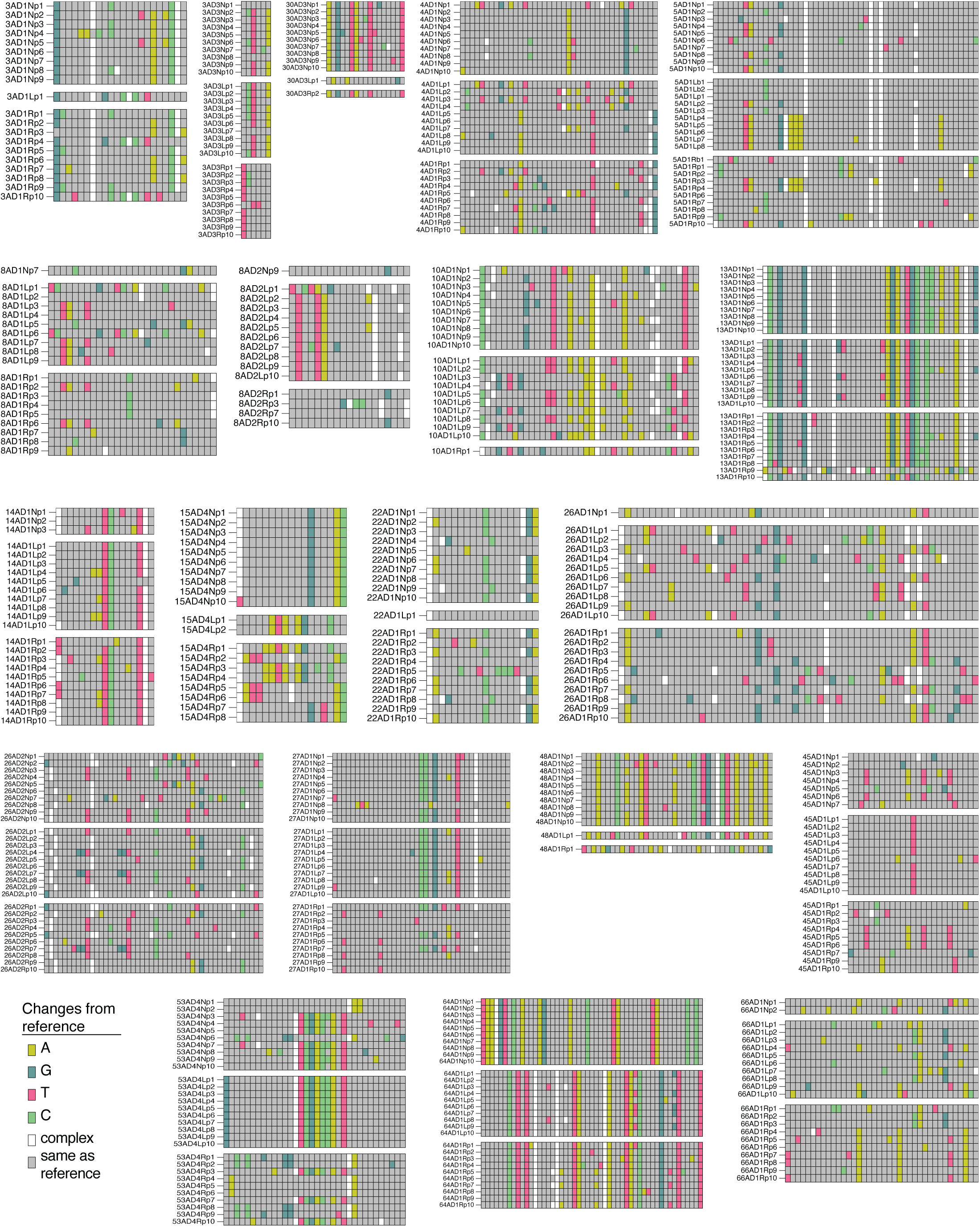
Heat maps of sites of nucleotide variants of analyzed isolates from all included subjects. Refers to **Fig. 4e, f**. Subjects were selected that had at least 1 isolate sequenced per site and 10 isolates sequenced total within a single timepoint. Each row represents an isolate, and each column represents a site of nucleotide variation. Sites consistent amongst all isolates are masked. Isolates with “N”, “L”, and, “R” in the row name were isolated from the nares, lesional skin, and rectum respectively.

**Extended Data Fig. 6:**
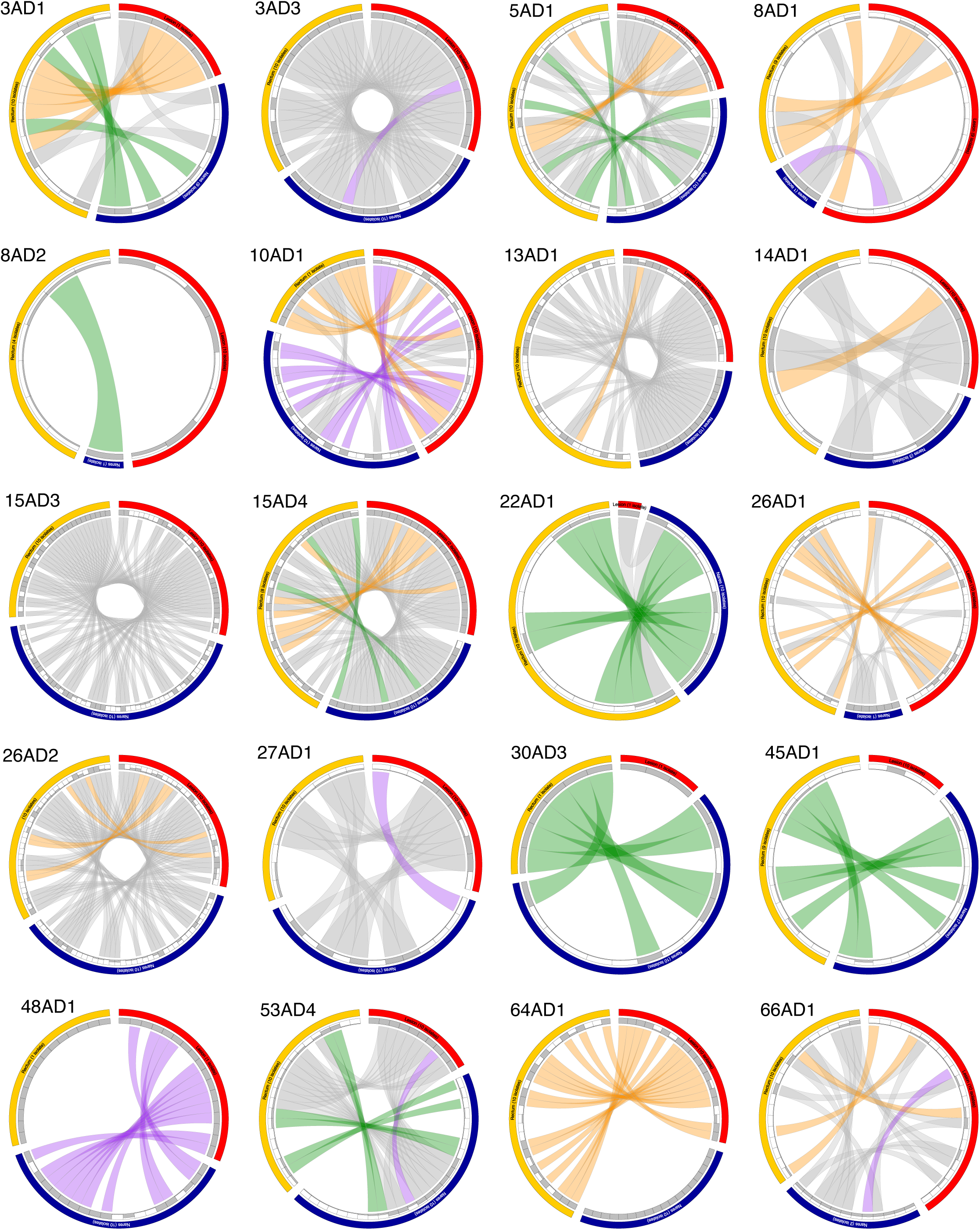
Shared nucleotide variants by body site of analyzed isolates from all included subjects. Refers to **Fig. 4e, f**. Subjects were selected that had at least 1 isolate sequenced per site and 10 isolates sequenced total within a single timepoint. Circular plots demonstrate uniquely shared variants across isolates from different sites. Outer arcs represent isolates from specific sites, while inner arcs show the proportion of isolates with variant at each site (bar chart). Ribbons connect sites that share variant: Gray ribbons show shared variants shared between all three body sites, and colored ribbons show those uniquely shared between two body sites.

**Extended Data Fig. 7:**
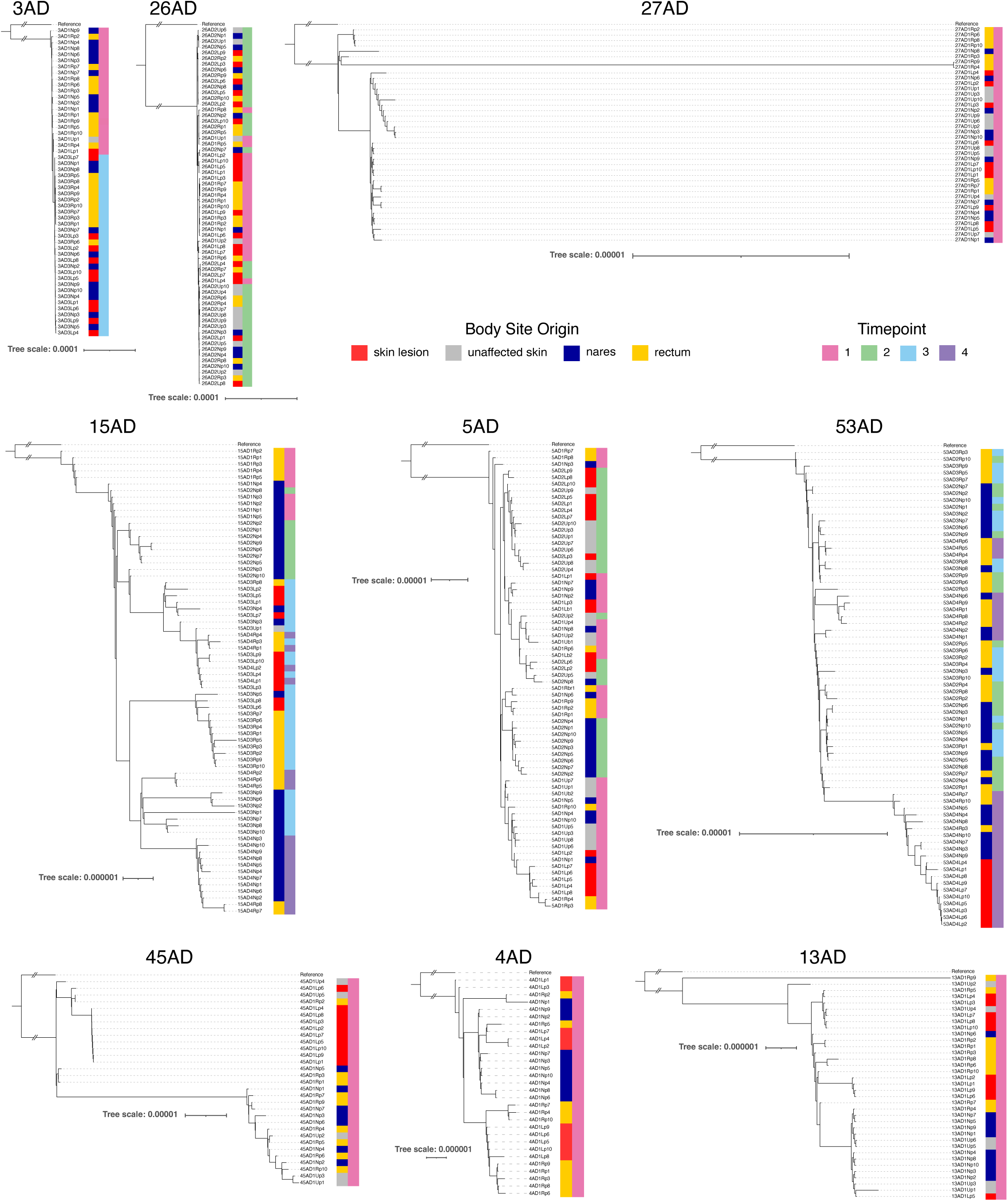
Maximum likelihood phylogenies of *S. aureus* isolates from subjects included in *S. aureus* genome diversity by body site analysis. Refers to **Fig. 4i**. Subjects were selected that had at least 7 isolates sequenced per site for all body sites within a single timepoint. Note, only the isolates from the timepoint for which 7 isolates sequenced per site for all body sites were available were included in Fig. 4i. However, all sequenced isolates per subject are shown here (regardless of timepoint). The reference genome for each phylogeny is the RefSeq genome assembly with the highest average nucleotide identity for the *S. aureus* lineage isolated from the individual subject (see **Methods**).

**Extended Data Fig. 8:**
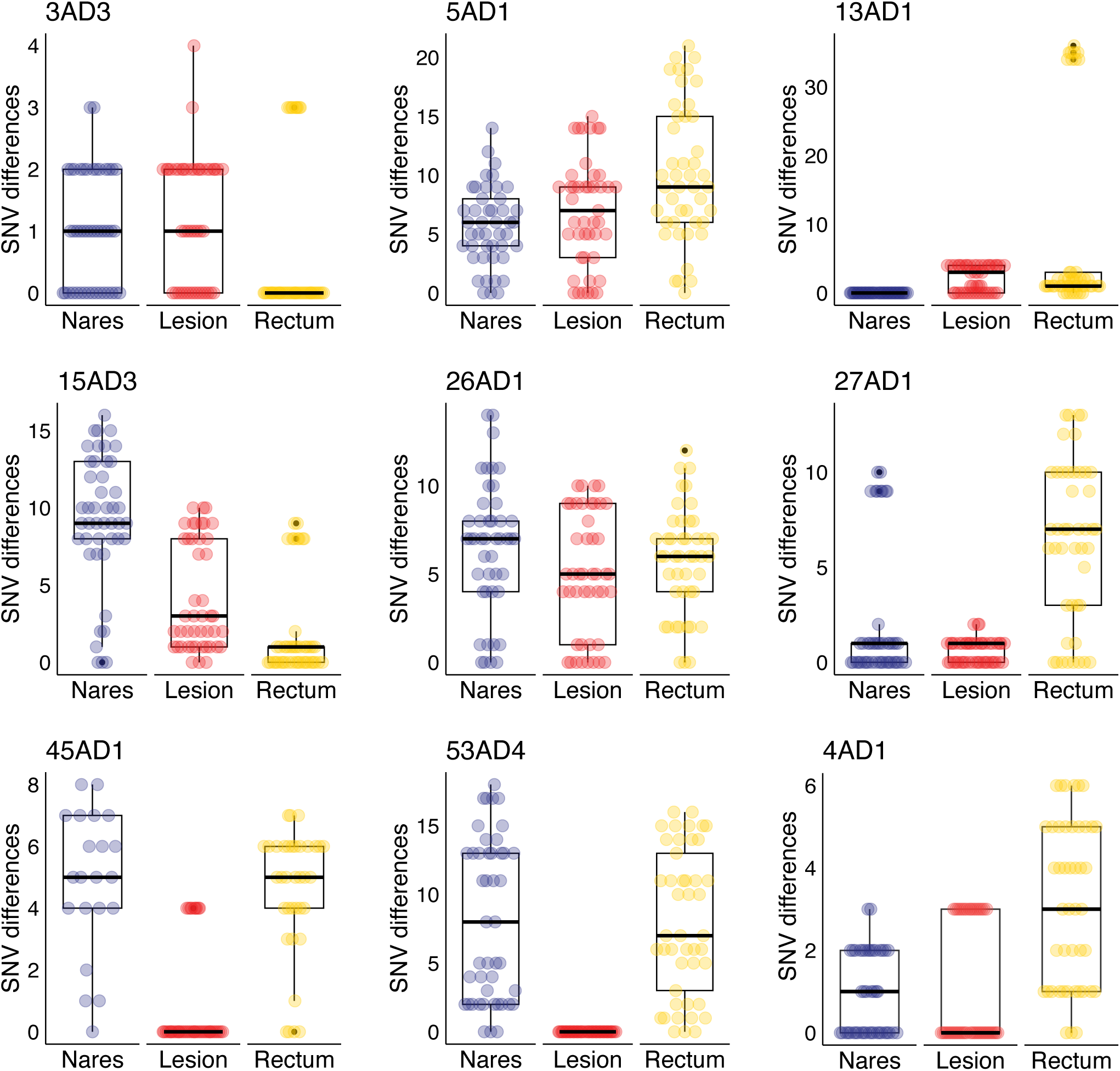
Within body site *S. aureus* genomic diversity. Refers to Fig. 4i. Subjects were selected that had at least 7 isolates sequenced per site for all body sites within a single timepoint. Note, only the isolates from the timepoint that fulfilled the criterion of at least 7 isolates sequenced per site for all body sites were included. Boxplots showing pairwise differences in number of single nucleotide variants (SNVs) between otherwise isogenic isolates from different body sites within an individual subject. Boxplot centered around median, lower and upper hinges indicating 1^st^ to 3^rd^ interquartile range (IQR), and whiskers indicating observations +/- 1.5 * IQR.

**Extended Data Fig. 9:**
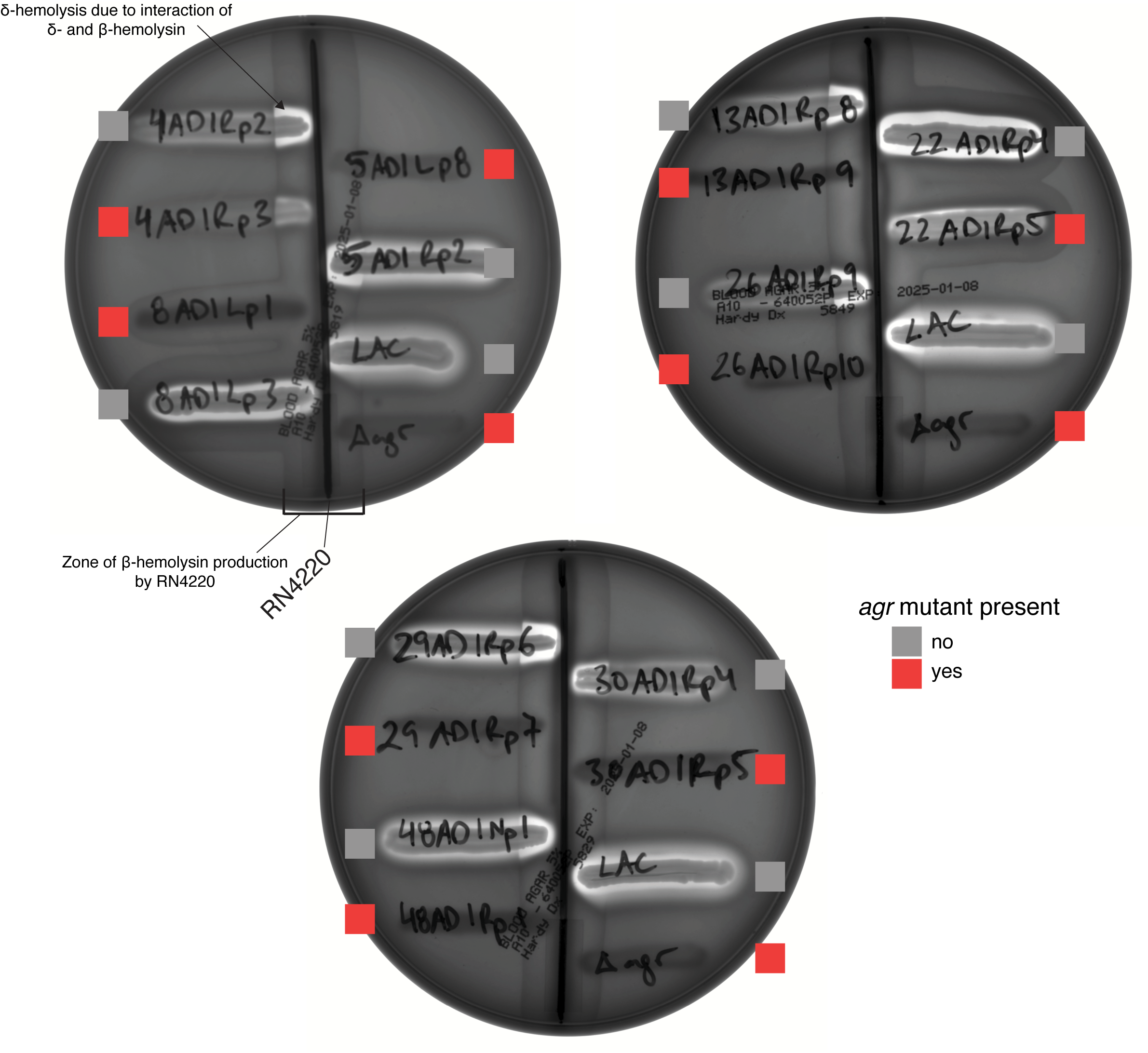
*S. aureus* isolates with *agr* mutations have decreased δ-hemolysin production. Images of three TSA plates with 5% sheep’s blood with labelled isolates cross-streaked to β-overproducing strain RN4220.^40^ Selected isolates are *agr*-WT vs. *agr*-mutated isolate pairs from the same lineage. LAC is shown as positive control and LACΔ*agr* as negative control. δ-hemolysis observed as zone of clearing in area of interaction of δ- and β-hemolysin.

**Extended Data Fig. 10:**
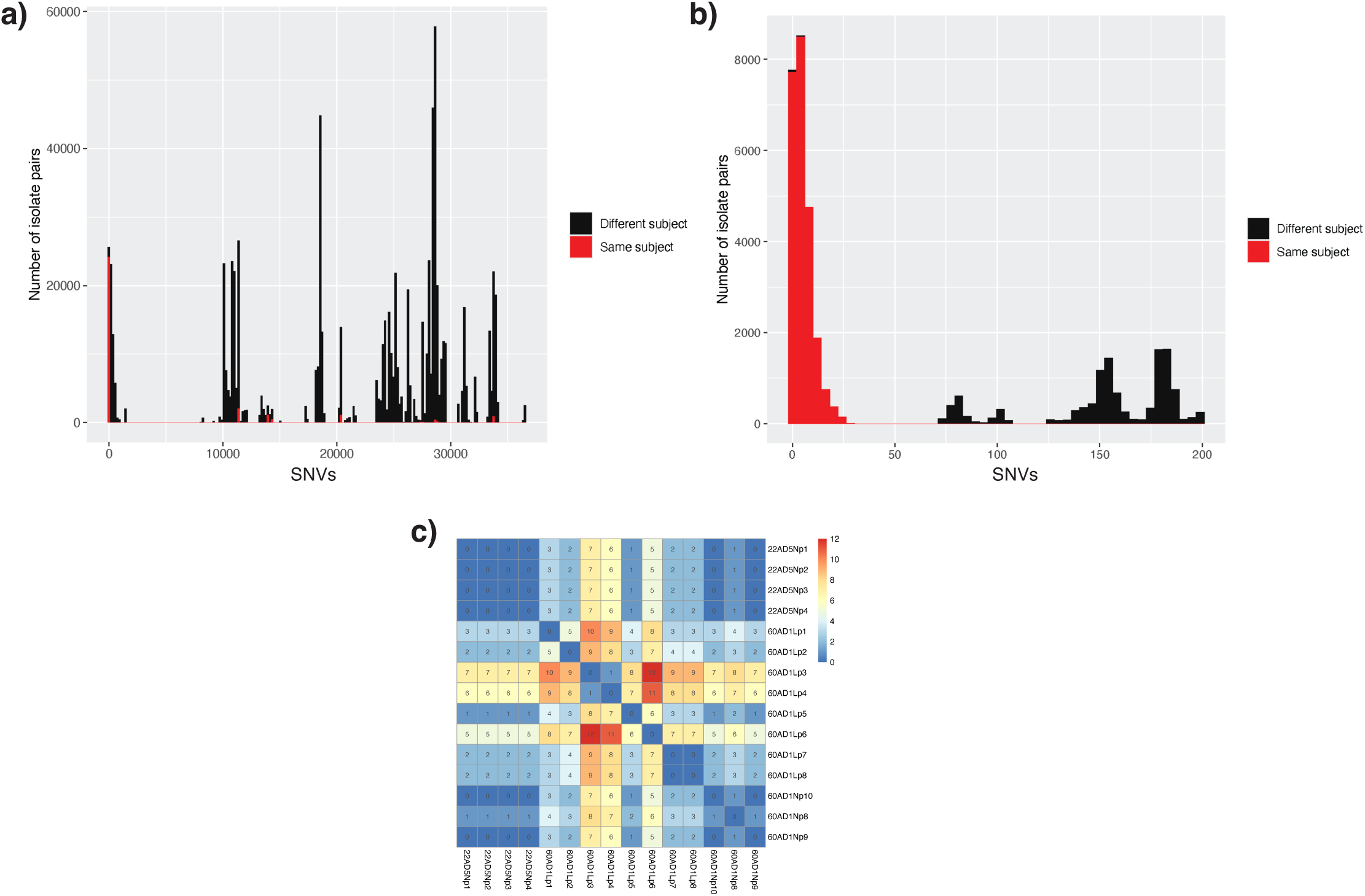
*S. aureus* isolates related by 50 single nucleotide variants or less predominantly are obtained from the same individual. **a)** Number of single nucleotide variant (SNV) differences between two isolates in AD cohort. Isolate pairs labelled by whether they were derived from the same subject or two different subjects **b)** Magnified aspect of prior graph showing isolate pairs with 200 SNV difference or less. **c)** SNV matrix of isolates from subjects 22AD and 60AD that clustered together using a 50 SNV threshold.

## Notes

### Competing Interest Statement

The authors have declared no competing interest.

